# Infection-Specific Reprogramming of Microglia Reveals Distinct Virulence Pathways Linking Periodontal Pathogens to Alzheimer’s Disease

**DOI:** 10.1101/2025.06.06.658266

**Authors:** Marta Kaminska, Noemie A. Dudzinska, Anne Klapper, Stefanie Geissler, Nadine Taudte, Sebastian Greiser, Jan Potempa, Holger Cynis, Piotr Mydel

**Affiliations:** Broegelmann Research Laboratory, Department of Clinical Science, Faculty of Medicine, University of Bergen, Bergen, Norway (;;); Fraunhofer Institute for Cell Therapy and Immunology, Department of Molecular Drug Design and Target Validation, 06120 Halle (Saale), Germany (;;); Periotrap Pharmaceuticals GmbH, Weinbergweg 22, 06120 Halle, Germany; Fraunhofer Institute for Cell Therapy and Immunology, Perlickstraße 1, 04103 Leipzig, Germany; Department of Microbiology, Faculty of Biochemistry, Biophysics and Biotechnology, Jagiellonian University, Kraków, Poland; Department of Oral Immunity & Infectious Diseases, University of Louisville School of Dentistry, Louisville KY, USA; Junior Research Group “Immunomodulation in Pathophysiological Processes” Faculty of Medicine, Martin Luther University Halle-Wittenberg, Germany

## Abstract

Microglial dysregulation is increasingly recognized as a driver of Alzheimer’s disease (AD), yet how pathogen-specific cues sculpt microglial diversity remains unclear. Here we integrate high-dimensional single-cell cytometry in vitro with spatial proteomics in vivo to dissect the impact of two major periodontal pathogens on microglia. Using a 36-marker CyTOF panel, we exposed SIM-A9 microglia to wild-type Porphyromonas gingivalis (Pg) or Tannerella forsythia (Tf) and to gingipain-deficient or S-layer–deficient mutants, resolving 38 clusters. Virulence-factor “switches” redirected cells from homeostatic states toward i) oxidative, antigen-presenting programmes driven by Pg gingipains and ii) an immunosuppressive, exhausted-like state driven by the Tf S-layer. Complementary 37-marker imaging mass cytometry of 5xFAD × hTau knock-in mice chronically infected with Pg identified 21 microglial subclusters. The cortex—but not hippocampus—lost two Arg1⁺/IL-10⁺ immunoregulatory subsets (>2-fold decrease) while NADPH-oxidase-high microglia accumulated around amyloid-β and tau aggregates. These data demonstrate pathogen-specific reprogramming of microglia across model systems and brain regions, linking virulence-factor activity to AD-relevant neuroinflammation. By pinpointing gingipains and the bacterial S-layer as molecular “switches,” our study highlights tractable therapeutic targets for limiting infection-driven microglial dysfunction in Alzheimer’s disease.

## INTRODUCTION

Neuroinflammation is a hallmark of neurodegenerative diseases, including Alzheimer’s disease (AD), the most prevalent cause of dementia worldwide^1^. While inflammatory cascades in the central nervous system (CNS) were once considered secondary to neuronal loss, growing evidence now places microglia at the forefront of driving and modulating AD progression^2^. Under homeostatic conditions, these resident immune cells of the CNS support neuronal function by clearing debris and maintaining synaptic integrity. However, in AD, microglia can adopt disease-associated (DAM) states marked by excessive production of proinflammatory mediators, impaired phagocytosis, and enhanced neurotoxicity^2^.

Recent single-cell and single-nucleus RNA sequencing have revealed unexpected heterogeneity within microglial populations^3,4^. Specific marker signatures distinguish these subpopulations: for instance, up-regulation of CD86 and MHC II^5,6^ highlights increased antigen presentation, whereas PD-L1 (CD274) often reflects attempts to regulate T cell responses^7^. Concurrently, microglia expressing CD40 can amplify co-stimulatory signals that exacerbate neuroinflammation^8^. In contrast, TGFβ and IL-10 are frequently associated with immunoregulatory or “resting” phenotypes that safeguard neurons from inflammatory damage^9^. Enzymes such as NADPH oxidase further amplify reactive oxygen species (ROS) generation, linking chronic oxidative stress to neurodegenerative cascades^10,11^. Dissecting how these key markers change in pathological conditions is central to understanding - and potentially mitigating - the microglial contribution to AD pathogenesis.

Although multiple factors regulate these microglial transitions, a growing body of evidence implicates infectious agents as potent drivers of chronic neuroinflammation. The periodontal pathogen *Porphyromonas gingivalis* (*Pg*) has garnered special attention, given that its cysteine proteases (gingipains) have been detected in AD brains and shown to trigger neurodegeneration in animal models^12–14^. Another prominent periodontopathogen, *Tannerella forsythia* (Tf), carries a semicrystalline S-layer integral to bacterial adhesion and immune evasion and secretes an array of proteolytic enzymes potentially contributing to inflammation^15,16^. A direct impact of these virulence factors on microglial phenotypes remains poorly understood. Both organisms are implicated in periodontitis, and epidemiological and interventional studies suggest that they may contribute to cognitive decline and neuroinflammatory processes^17–19^. Periodontal disease frequently involves polymicrobial biofilms that include *Treponema denticola* or *Fusobacterium nucleatum*^20–22^ and may even involve co-infection with fungi (*Candida albicans*) or viruses (HSV-1, EBV), expanding the range of microbial signatures that microglia confront^23–26^. Polymicrobial synergy can converge on shared TLR or inflammasome pathways, amplifying neuroinflammatory circuits that might predispose to AD^27–29^. Since different virulence factors likely engage distinct microglial signalling pathways - some fuelling inflammation while others skewing microglia toward immunoregulatory or “exhausted” states - elucidating these pathogen-specific effects is critical for clarifying how oral infections might influence AD progression. However, robust mechanistic data at the protein level - detailing precisely how each pathogen reshapes microglial marker expression over time - remain sparse. While transcriptomic approaches have revolutionized our understanding of microglial diversity^30^, complementary high-dimensional proteomic analyses are essential for clarifying functional phenotypes and virulence factor– driven reprogramming.

In this study, we employ a 36-marker panel in cytometry by time-of-flight (CyTOF) to interrogate the phenotypic breadth of microglia exposed to *Pg* and *Tf*. By analysing wild-type strains alongside isogenic mutants (gingipain-deficient *Pg* and S-layer–deficient *Tf*), we identify virulence factor-specific pathways that shift microglia from resting or homeostatic states toward oxidative, antigen-presenting, or immunoregulatory phenotypes. We then extend these insights *in vivo*, performing spatial evaluation of microglial subpopulations in a 5xFAD x hTau-knock-in mouse model orally infected with *Pg*, using a high-dimensional imaging mass cytometry platform. Taken together, our results reveal a *Pg-*driven depletion of moderately activated microglia in favour of strong proinflammatory responses, offering compelling evidence for a contributory mechanistic link between chronic periodontitis and AD pathology - and underscoring the therapeutic and/or preventive potential of targeting specific bacterial virulence factors (e.g., gingipains) to preserve microglial homeostasis in the aging brain.

## RESULTS

### High-Dimensional CyTOF Profiling Reveals Extensive Microglial Heterogeneity

We employed a high-dimensional single-cell approach *via* CyTOF with a 36-marker panel to capture the breadth of microglial phenotypes following exposure to *Pg* and *Tf* at two timepoints (24h and 48h) and two MOIs (5 and 10) using the SIM A9 microglia cell line^31^. All used antibodies were optimised and titrated in-house, with target selectivity confirmed by staining SIM A9 stimulated with IFNγ. After de-barcoding and data pre- processing, which included gating out cell debris and beads, we downsampled all unique biological replicates to equal cell numbers.

Control (uninfected) SIM-A9 microglia primarily expressed “resting” markers (TGFβ, IL-10, CD206, Arg1), consistent with a homeostatic or surveillant phenotype in the absence of acute inflammatory stimuli (Fig. 1). Upon infection, these resting phenotypes rapidly diverged, exhibiting pathogen-specific patterns of activation, suppression, or phenotypic exhaustion. *Pg* triggered robust upregulation of proinflammatory (TNFα, IL-12p40) and oxidative markers (NADPH oxidase, iNOS), often peaking around 24 hours, while *Tf* induced a more immediate global suppression of major activation molecules (CD40, CD44, MHCII, NADPH oxidase) in certain clusters, hinting at a distinct immunomodulatory or “exhausted” route (Fig. 1).

**Figure 1.**
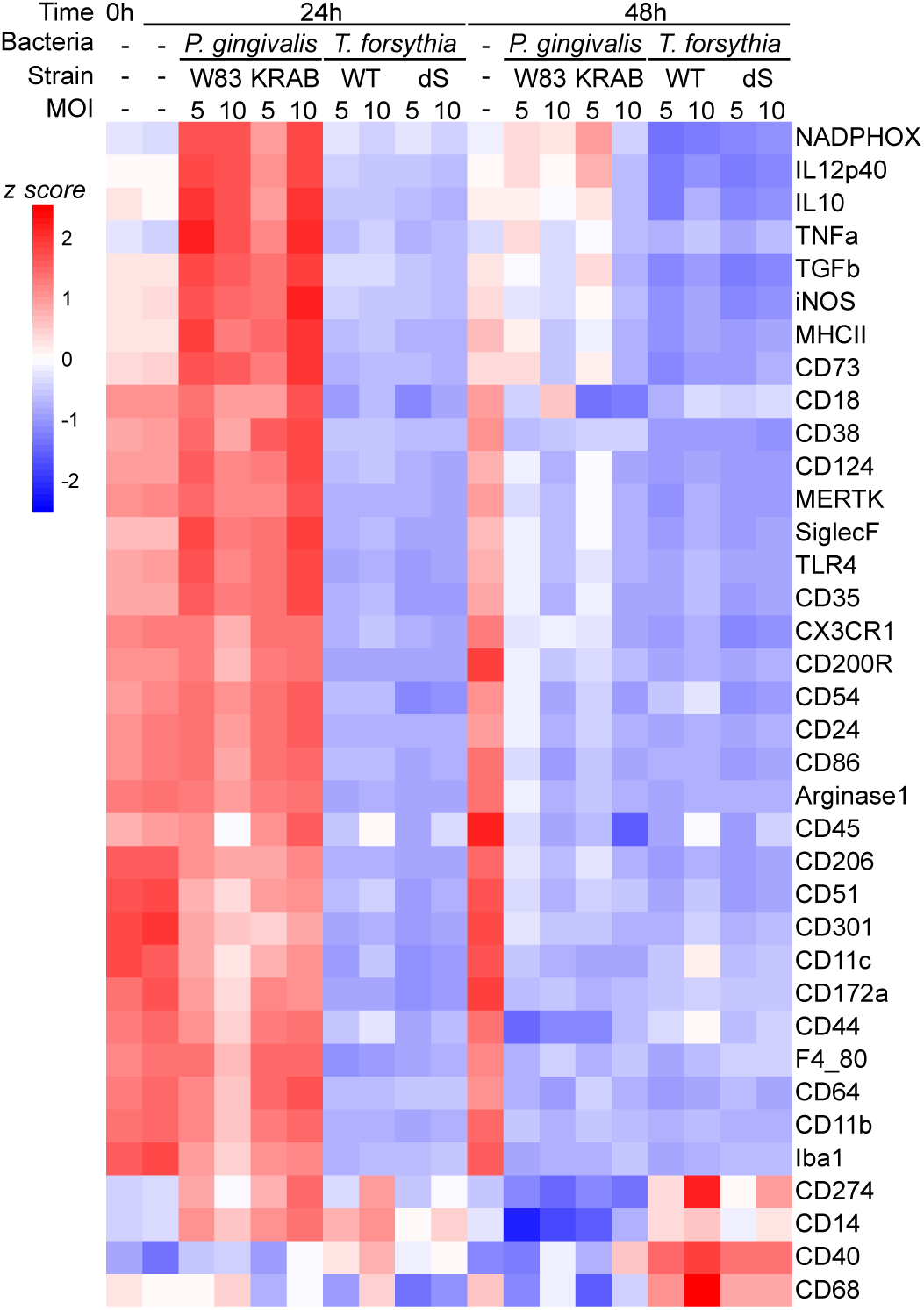
SIM-A9 microglia mount a rapid inflammatory burst to *P. gingivalis* and acquire an exhaustion-/senescence-like profile after *T. forsythia* infection. Heat map of mean arcsinh-transformed, z-scored intensities for 36 CyTOF-measured protein markers in SIM-A9 microglia that were left untreated (0 h) or infected for 24 h and 48 h with either *P. gingivalis* (Pg; W83 or KRAB) or *T. forsythia* (Tf; WT or dS) at multiplicities of infection (MOI) 5 or 10. Each column is the average of all single-cell events pooled from three independent cultures per condition (745,000 cells after down-sampling); rows are hierarchically clustered. Pro-inflammatory markers (for example, TNF, NADPHOX) peak 24 h after *Pg* exposure, whereas exhaustion/senescence markers (for example, PD-L1) are preferentially up-regulated following *Tf* infection. Data are z-scored per channel across biological replicates. See Supplementary Table 1 for the antibody panel and Supplementary Fig. 1 for gating strategy.

### Rapid Divergence from Homeostatic Baselines

By using R-based Phenoannoy, we have identified 38 distinct microglial clusters, each distinguished by unique constellations of inflammatory, oxidative, antigen-presenting, and homeostatic markers (Fig. 2). In control conditions, over 60% of total microglia aggregated into clusters highly expressing the hallmark homeostatic triad (TGFβ, IL-10, CD206; Fig 2B and Extended Data Fig. 1A). Only ∼10% of cells fell into mild “activation-leaning” states. Upon pathogen exposure, these resting clusters diminished—often by >70% at 24 hours (p < 0.01)—alongside expansions of more activated or exhausted subsets. Specifically, *Pg* infection drove marked elevations in TNFα, IL-12p40, MHCII, and iNOS, whereas *Tf* promoted broad marker downregulation, suggesting an alternative immunosuppressive or senescent microglial pathway (Fig. 2A-B). To confirm these cluster-level observations, we conducted quantitative comparisons of cluster abundance across biological replicates (Fig. 2B). Scatter plots for key clusters (Extended Data Fig. 1B) indicated significantly higher abundance enrichment under *Pg* wild-type (*Pg* WT) at MOI=10 for clusters 22, 23, 27, 35 versus control (p < 0.05 or p < 0.01) or *Tf*-infected versus control for cluster 10 (p < 0.05 or p < 0.01). Conversely, virulence-deficient mutant exposures (gingipain-deficient *Pg* KRAB, or strain *Tf* dS not producing S-layer) often yielded similar tendencies but at a reduced scale, thus suggesting full virulence is necessary for maximal proinflammatory or immunomodulatory reprogramming.

**Figure 2.**
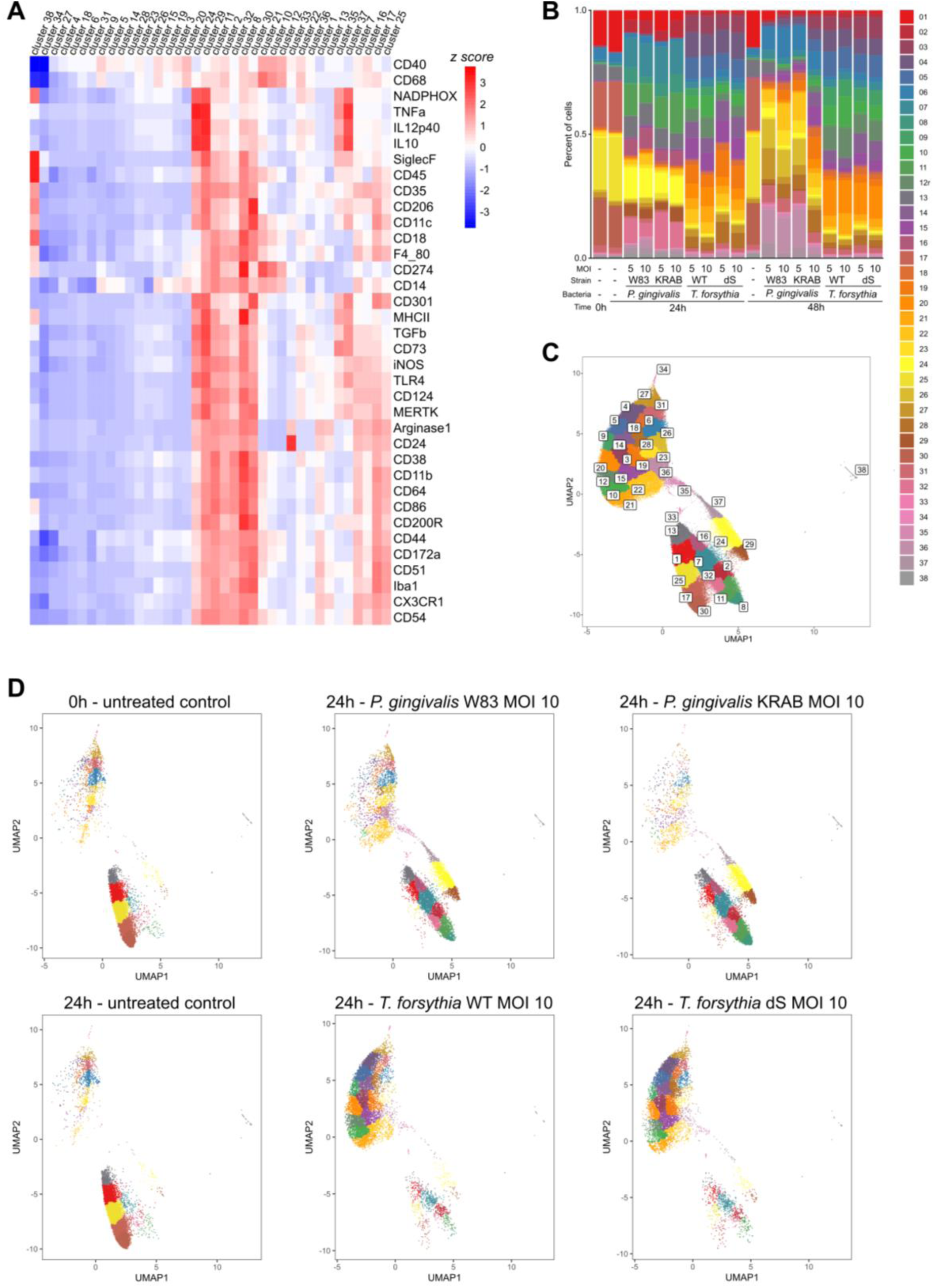
SIM-A9 microglia adopt discrete phenotypic states after infection with *P. gingivalis* or *T. forsythia*. **A.** Heat map of z-scored, arcsinh-transformed intensities for 36 CyTOF-measured proteins across the 14 clusters identified by Phennoannoy (k = 800) from the pooled dataset of 745,000 single cells (three independent cultures per condition; see Fig. 1 and Methods). Rows are markers, columns are clusters ordered by hierarchical clustering. **B.** Relative abundance of each cluster (percentage of total cells) in untreated microglia (0 h) and after infection with *P. gingivalis* strains W83 or KRAB or *T. forsythia* strains WT or S-layer deficient (Tf dS) at multiplicities of infection (MOI) 5 or 10 for 24 h and 48 h (bars, mean ± s.d.; n = 3 biological replicates). Differences among conditions were assessed with a Kruskal–Wallis test (P < 0.0001) followed by Dunn’s multiple-comparison correction; filled symbols indicate Padj < 0.05. **C.** UMAP projection of the concatenated dataset, coloured by cluster identity. **D.** Representative UMAPs for untreated cells (0 h) and for each MOI 10 pathogen condition at 24 h, coloured as in c, illustrate pathogen-specific redistribution of clusters. Data derive from the same infections and CyTOF acquisition described in Fig. 1; gating, down-sampling and clustering parameters are detailed in Methods.

### Time-Dependent Dynamics of Microglia upon Exposure to Periodontopathogens

Analyses at 0, 24, and 48 hours for each experimental condition (*Pg* WT, *Pg* KRAB, *Tf* WT, *Tf* dS; MOIs 5 or 10) revealed notable heterogeneity in how cluster abundances changed over time in response to infection. Three clear patterns have emerged: (***i)*** several clusters spiked at 24 hours before returning near baseline by 48 hours (e.g., cluster 24 with high NADPH oxidase, TNFα, IL-12). This “flash and burn” motif was prominent in response to *Pg* WT (Fig. 2B, Extended Data Fig. 1C) and, to a lesser extent, in *Tf* WT infections (Fig. 2B, Extended Data Fig. 1D). For example, cluster 15, which showed CD14^HIGH^ and CD68^HIGH^ expression, peaked specifically in *Tf*-infected group at 24 hours and then was markedly reduced in abundance by 48 hours, illustrating an ephemeral immunomodulatory state that may later transition to an exhausted profile if pathogen exposure continues. ***(ii)*** some clusters with populations negligible at 24 hours, surged significantly at 48 hours. These late abundance clusters such as 26 and 31 —generally low in CD14/CD40—rebounded in response to infection with *Pg* KRAB or *Tf* dS, suggesting a delayed or compensatory microglial activation wave when virulence factors are absent or less potent (Fig. 2B, Extended Data Fig. 2A). ***(iii)*** finally, the abundance of cells in Arg1^HIGH^ or CD206^HIGH^ clusters (e.g., clusters 17, 25, 30; Fig. 2B and Extended Data Fig. 1A) decreased in response to all pathogen challenges, regardless of the expression of virulence factors. This consistent decline argues that these homeostatic microglia are not phenotypically preserved in the face of proinflammatory or immunomodulatory stressors.

### *P. gingivalis* Wild Type Drives Strong Proinflammatory Polarization

In *Pg*–infected SIM A9 cultures, we observed high expression of canonical inflammatory and antigen-presenting markers - CD86, MHC II, TLR4, NADPH oxidase, iNOS—within certain subpopulations that peaked around 24 hours (Extended Data Fig. 1C). Clusters 24 and 29, for instance, showed elevated TNFα and IL-12p40 compared to control levels (p < 0.01). Comparisons at MOI 5 vs. 10 showed that higher bacterial load elicited stronger cluster expansions. By 24 hours under MOI 10, these “proinflammatory peak” clusters 24 and 29 more than doubled in proportion (Extended Data Fig. 1C). Interestingly, by 48 hours, many of these clusters returned to near- or sub-baseline expression, suggesting partial immune resolution or cellular exhaustion. Such biphasic responses likely involve initial microglial recognition of bacterial ligands (e.g., *via* TLR2/4), prompting acute cytokine and ROS release. In some subsets, including cluster 22, we detected iNOS co-expression with CD38, pointing to a state of sustained oxidative capacity with a NAD+ metabolic shift that could impact neuronal health if it persists beyond 48 hours. These dynamics emphasizes the importance of analysing multiple time points and infection doses to capture microglial complexity.

### Role of Gingipains in Shaping Microglial Responses

Comparative studies using gingipain-deficient *Pg* KRAB consistently highlighted gingipains as central drivers of the strong inflammatory shift in the infected microglia (clusters 6, 16, 23, 24, 35, 37, Fig. 2B, Extended Data Fig. 1B-C and 2B). While *Pg* WT markedly promoted the expansion of clusters expressing high levels of NADPH oxidase, TLR4, iNOS, and IL-12p40, *Pg* KRAB-infected microglia showed more muted or delayed responses. Clusters maintaining higher CD206 or TGFβ were severely diminished by *Pg* WT but persisted under *Pg* KRAB, implying that gingipains help reprogram microglia from a regulatory/homeostatic phenotype to an inflammation-prone or antigen-presenting state. These data resonate with reports linking gingipains to exacerbated neurotoxicity and microglial hyperactivation^32,33^, suggesting that targeting gingipains might preserve beneficial microglial subsets—an important consideration for halting pathogen-driven neurodegenerative cascades^34^.

### S-Layer in *T. forsythia* is a Key Inflammatory Driver

In comparison to *Pg*, *Tf* elicited more moderate oxidative/proinflammatory shifts, but induced mild antigen-presentation phenotype (CD40, CD14, CD86) in specific clusters (i.e., 10, 12, Extended Data Fig. 1B and 2C), suggesting a different route to microglial engagement. Many *Tf*-induced changes were statistically lower in magnitude than robust expansions triggered by *Pg* (p < 0.05). Deleting the *Tf* S-layer (strain *Tf dS*) further reduced these moderate effects, dropping cluster abundance close to control. This aligns with the established importance of the S-layer as a major virulence factor for *Tf*. The absence of the S-layeraccelerated emergence of senescent clusters lacking typical activation markers (CD14, CD40). Hence, while *Tf* has not yet been studied in the context of the AD pathogenesis, in can be assumed that its repeated dissemination into the brain can cause low-level microglial disturbance contributing to chronic CNS inflammation. Moreover, in cluster 10 - uniquely upregulated under *Tf* WT (Fig. 2B and Extended Data Fig. 1B) - the co-expression of PD-L1, CD14 and CD68 suggested a subtle immunomodulatory axis that might alter T-cell interactions if sustained over longer time frames (Extended Data Fig. 1B). Extending infection times (beyond 48h) or applying co-infection models may help clarify *Tf*’s longer-term significance.

### Convergence with Neuroinflammatory Phenotypes

Although these in vitro data alone do not confirm the AD pathology, some cluster-level marker profiles—such as iNOS^HIGH^, IL-12^HIGH^, CD64^HIGH^, or CD86^HIGH^ - overlap with neuroinflammatory or disease-associated microglia (DAM) reported in AD contexts. For instance, cluster 37, which peaked at 48 hours under *Pg* WT, aligned closely with heightened inflammatory states seen in late-stage neurodegeneration (Extended Data Fig. 2B). Our results imply that repeated or chronic exposure to periodontal pathogens could prime microglia toward the AD-like phenotypes if these inflammatory states remain unresolved.

### Multilayered Functional States: Illustrative Cluster Examples

Despite highly rigorous *in vitro* conditions, SIM A9 displayed striking heterogeneity and diverse functionality in response to pathogens and their unique patterns of virulence factors. For example, cluster 10 (CD40^HIGH^, CD68^HIGH^, PD-L1^HIGH^, Extended Data Fig. 1B) was markedly upregulated after *Tf* WT infection, but completely absent in presence of *Pg*. The S-layer deletion reduced its abundance, implying the S-layer fosters immunoregulatory or checkpoint-rich microglia. Similarly, gingipain-induced cluster 24 (NADPHox^HIGH^, IL- 12H^HIGH^, IL-10^MOD^, TLR4^MOD^) typically exhibited highest abundance at 24 hours then waned by 48h. Cluster 35 (CD73^HIGH^) was remarkably time-dependent, peaking at 48h in presence of *Pg* KRAB, hinting at adenosine-mediated immunosuppression emerging in the absence of gingipain-driven hyperinflammation^35^. Lastly, cluster 28 (CD14^LOW^, CD40^LOW^, Extended Data Fig. 1D) was observed in both *Pg* and *Tf* infections, but earlier in *Tf* and at higher abundance in the absence of S-layer, suggesting pathogen-specific timing for “exhausted/senescent” microglial states.

### 5xFAD x hTau KI Mice Mirrors Key In Vitro Findings

Overall, our *in vitro* CyTOF data reveal that *Pg -* particularly *via* gingipains - drives potent proinflammatory or antigen-presenting microglial states, whereas *Tf*’s S-layer fosters subtler but still disruptive immunomodulatory or exhausted phenotypes. While these findings illuminate how periodontal pathogens exploit virulence factors to reshape microglial states *in vitro*, caution must be applied when extrapolating to the clinical AD setting. Immortalized lines like SIM A9 cannot fully model the interactive *milieu* of the aging brain, which includes astroglial crosstalk, neuronal signals, and peripheral immune involvement. Hence, we pursued extended *in vivo* experiments in 5xFAD mice crossed with human Tau knock-in mice-The mice generate high amyloid levels and showing pathological conformations of tau. To determine whether the pathogen-induced microglial shifts observed *in vitro* extend to an AD-like context, the recently published 5xFAD x hTau KI mice^36^ underwent a 22-week oral infection protocol with *Pg* wild type (or mock PBS). Mice were sacrificed at ∼11.5 months and brains were collected, and sectioned. To facilitate comparisons between experimental groups, two brain sections per group were placed onto single slide, at a total of 4 (vehicle) or 3 brain sections (*Pg*-infected) were scanned. We analyzed sections *via* imaging mass cytometry using a 38-antibody panel overlapping with, but extending beyond, the *in vitro* set to include markers of tissue architecture (GFAP, NeuN, Cox2, GLUT1, CD13, Vcam1) that were used to identify cell types and AD lesions (amyloid, Tau, pTau). Cell-like objects were segmented (Ilastik–CellProfiler), then clustered with RPhenograph, maintaining a single-cell analytical pipeline analogous to our CyTOF approach in SIM A9 cultures.

### Regional Variations and Microglial Subset Composition

An initial characterization of segmented objects revealed region-specific differences in marker expression following infection, particularly between the cerebral cortex and hippocampus (Fig. 3A). The cortex exhibited higher baseline levels of CD40, CD13, CD172a, Arg1, and Tau, with modest increases in Iba1 and Vcam1 signals, whereas the hippocampus showed modest post-infection downregulation of GLUT1, Cox2, CD13, and CD172a. There were no significant differences between groups in total microglial frequency, microglial ROI coverage, or Iba1 antigen levels, as further confirmed by immunofluorescence imaging (Extended Data Fig. 3). Semi-supervised clustering of the Iba1^HIGH^ microglial population (Fig. 3B–E) identified 21 distinct subclusters, several of which corresponded to oxidative (NADPH oxidase^HIGH^), antigen-presenting (CD86, MHCII), or regulatory (IL-10, Arg1) phenotypes previously observed in vitro. Most subclusters were present at similar frequencies in infected and control groups (Fig. 3F–G), suggesting subtle compositional shifts rather than global expansion or depletion of microglia.

**Figure 3.**
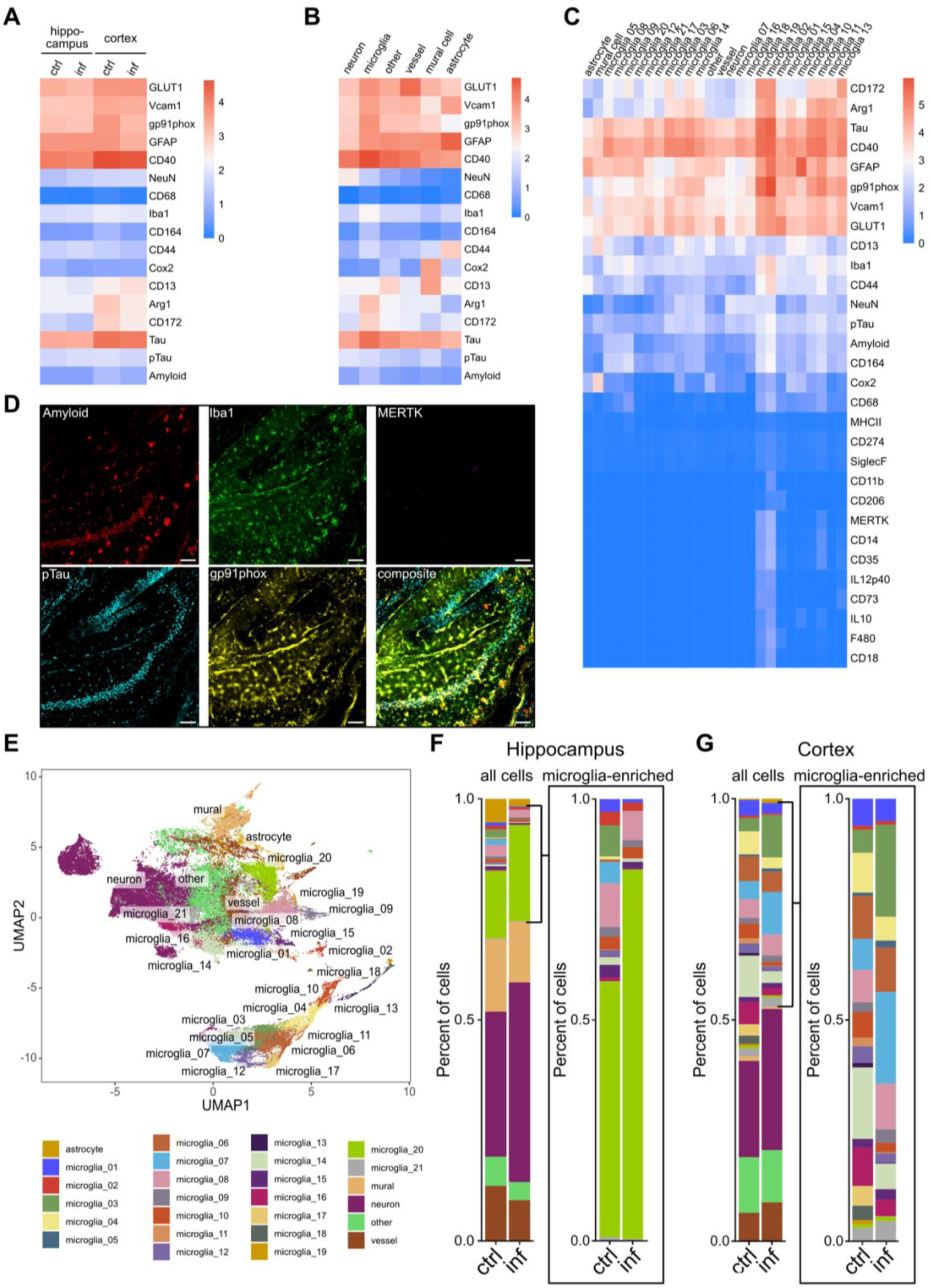
Region-specific microglial phenotypic shifts following chronic *P. gingivalis* exposure in hTau-knock-in × 5×FAD mice. **A.** Heat map showing arcsinh-transformed, z-scored expression of 17 imaging mass cytometry markers across all initial Rphenograph-derived clusters (k = 50; n = 78,622 cells). Columns represent individual imaging subgroups, and rows denote marker intensity. **B.** Marker expression heat map of broad cell classes identified by cluster-level profiling. Cell types were assigned based on characteristic marker patterns: NeuNHIGH (neurons), Iba1HIGH (microglia), GLUT1HIGH (endothelium), GFAPHIGH (astrocytes), CD13/COX2HIGH (mural cells), and ’other’ (unclassified). **C.** Heat map of arcsinh-transformed, z-scored marker expression across 21 microglial subclusters derived from secondary clustering (k = 60) of Iba1HIGH cells. **D**. Representative hippocampal image from the IMC dataset showing single-channel expression of Aβ (amyloid), Iba1, MERTK, pTau (AT8), and gp91phox, with composite overlay (far right). Scale bar, 100 µm. **E.** UMAP representation of batch-corrected events, coloured by the 21 subclusters defined in c, highlighting the phenotypic redistribution of microglia following infection. **F.** Relative abundance of microglial subclusters in hippocampus, displayed as proportion of total segmented events (mean per group). Group comparisons performed with Kruskal–Wallis test with Dunn’s post hoc correction; Padj < 0.05 were considered significant. **G.** As in F, but for cerebral cortex, revealing region-specific subcluster enrichment. Data are from PBS-treated control mice (n = 4) or *P. gingivalis*–infected mice (n = 3) after 22 weeks of oral challenge. Tissue processing, staining, denoising, segmentation, clustering, and statistical methods are described in detail in Methods.

However, the cortical microglial landscape showed a significant depletion of two subsets - microglia 10 and microglia 14 - in infected mice (Fig. 3G, p < 0.05, Kruskal–Wallis with Dunn’s post-hoc). Microglia 10 displayed a regulatory profile (relatively high Arg1, CD40, moderate IL-10, MERTK, CD14, CD35, MHCII), closely resembling the “mixed” or modestly activated subpopulations suppressed by *Pg in vitro*. Microglia 14, conversely, lacked IL-10, MERTK, or CD14 and showed low MHCII. Abundances of both were reduced more than two-fold in infected samples vs. controls, indicating a loss of immunomodulatory or mild-activation microglia in the cortex. In contrast, hippocampal microglia composition remained comparatively stable and arguably more homogenous (more than 50% of identified microglia belonged to subcluster 20) apart from moderate abundance increase in microglia 20 (CD172a^LOW^, Arg1^LOW^, CD40^MOD^), which partially parallels the “milder” or “exhausted” *in vitro* phenotypes observed under *Tf* exposure. These data indicate that hippocampal microglia possess region-specific thresholds for infection-driven reprogramming.

### Spatial Interactions and Protein Aggregate Association

To understand how these altered subpopulations might interact with the local environment, we examined cell-type–enriched regions (Fig. 4A-C) dominated by blood vessels (region 1), astrocytes (2), neurons (3), “other” cell types (4), mural cells (5), or microglial subclusters (6). Comparing their relative frequencies pre- vs. post-infection revealed that in the cortex, the ratio of “non-interacting” microglia (region 6 in spatial context 6 containing majority of the microglial subsets) decreased (from ∼0.35 ± 0.05 to ∼0.25 ± 0.04, p < 0.05), while spatial contexts (SCs) of neurons and microglia_21 interacting with microglia_08, _15 and “other” cell types (SC 3_4), but also with blood vessels and microglia_02 (SC 1_3_4) increased post Pg-infection (Fig. 4D). Clusters reminiscent of “flash-and-burn” microglia (e.g., microglia 8, 15, 21) became more prevalent in these multi-cell contexts, suggesting an increased immune engagement and cellular crosstalk post-infection. In parallel, microglial clusters 10 and 14, depleted during oral infection, were predominantly assigned to the non-interacting microglial context (SC 6, Fig. 4E). SC 4_6, harbouring microglia 14, was depleted in the cerebral cortex post-infection, similarly to SC 3_4_6 in hippocampus. In contrast to the changes observed for cortices, SC 3_4 was slightly depleted post-infection, similarly to SC 1, 1_6 (interaction between blood vessels+microglia_02 and microglia regions) and 5 (comprised of mural cells). Moreover, a hippocampal enrichment in SC 3 was observed post-infection, which was predominantly composed of neurons and approx. 50% of microglial subcluster 21, and to a smaller degree increase in abundance of SC 1_5 (blood vessels - mural cells) and 3_6 (neurons - microglia_20, the subcluster dominating in hippocampus). Although hippocampal changes were not as pronounced, the region harboured potential “compensatory” microglial subsets that appeared to resist infection-driven depletion.

**Figure 4.**
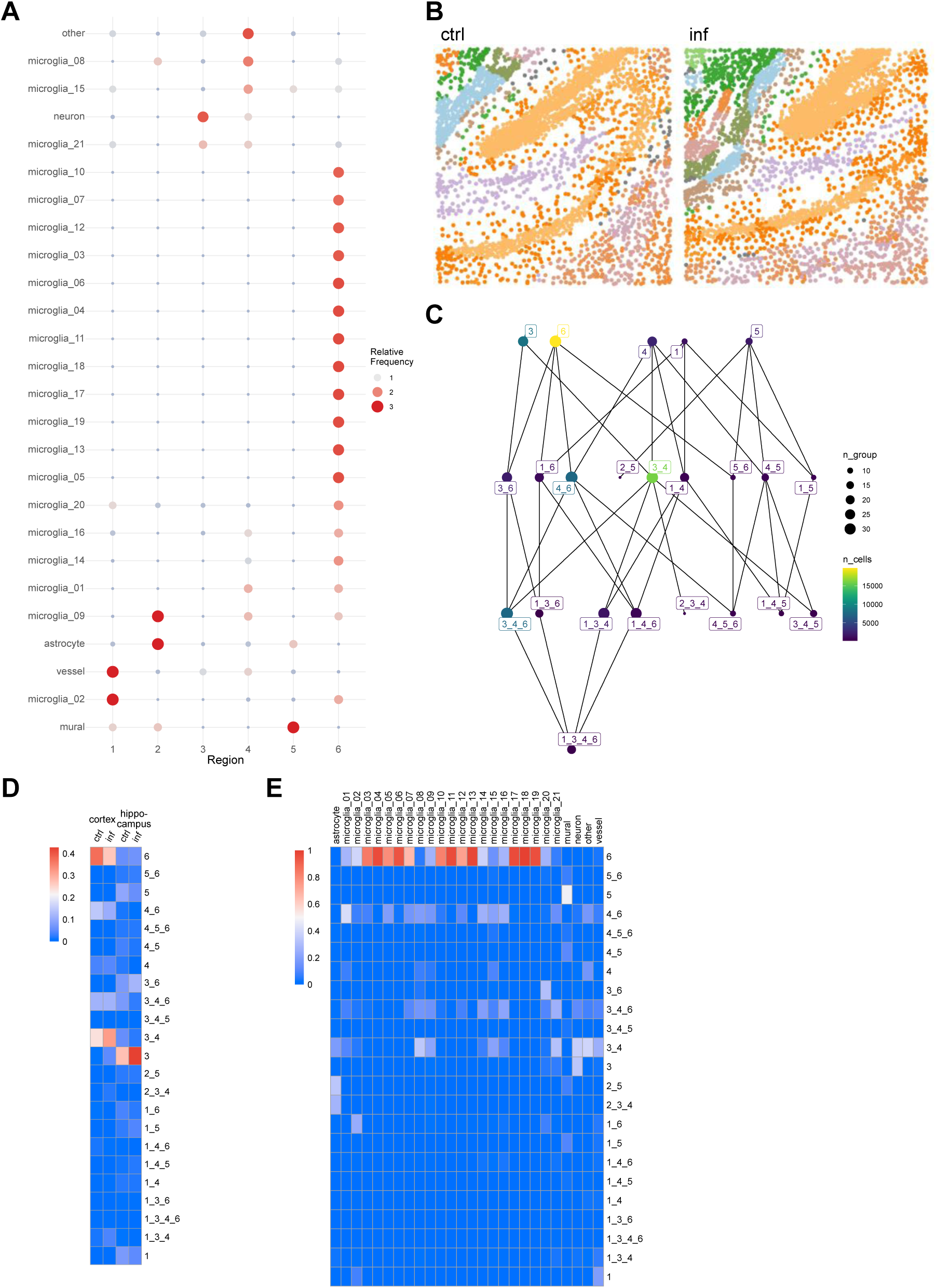
Chronic P. gingivalis infection alters spatial microenvironments in hTau-knock-in × 5×FAD mice. **A.** Segmented objects were assigned to one of six major cell-type–enriched regions (NeuNHIGH neurons, Iba1HIGH microglia, GFAPHIGH astrocytes, GLUT1HIGH endothelium, CD13/COX2HIGH mural cells, and undefined “other”) *via* expansion algorithm (radius = 40 px) based on full clustering (major cell types, with microglia further subclustered. **B.** Representative hippocampal ROIs from a PBS-treated control (left) and infected (right) mouse, with spatial contexts (SCs) shown in unique colors. SCs were defined using a k-nearest neighbor (k = 40) approach based on cellular proximity and composition. **C.** Overview of SC identities and their interrelationships, combining all retained ROIs (≥5 ROIs per SC, ≥200 total objects per SC). **D.** Frequencies of individual SCs per brain region and experimental group, shown as the proportion of all SCs within that category. **E.** Proportions of each identified cluster (21 microglial subclusters plus 5 remaing main cell types) within each SC, expressed relative to the total number of cells assigned to that SC. SCs were constructed by spatial graph modeling using imcRtools, followed by compositional and contextual analysis via lisaClust. Low-frequency SCs (occurring in <5 ROIs or with <200 cells total) were excluded from analysis.

To evaluate AD lesion interactions, we segmented the protein aggregates into four categories: (1) amyloid^HIGH^; (2) pTau (pTau^HIGH^ with NeuN^HIGH^); (3) pTau_d (pTau^HIGH^ but NeuN^LOW^); and (4) mixed (amyloid^HIGH^ and pTau^HIGH^) (Fig. 5A, D-E). Heatmap analyses of these lesions (23 chosen antigens) showed amyloids often induced robust Iba1 infiltration (implying strong microglial presence), while mixed lesions contained the highest gp91phox and CD40 signals, consistent with a potent oxidative or antigen-presenting environment. Proximity analyses (Fig. 5B–C) highlighted the microglia subset 10 as infiltrating the pTau- or amyloid-rich regions, with subset 14 more frequently observed at the lesion border. We then further performed proximity interaction analysis, which confirmed these observations: subset 10 exhibited highest interaction score with mixed-type lesions of all microglial subsets, pointing to a critical role of this subset in lesion formation or maintenance (Fig. 5F). Notably, microglia 10 and 14 - both diminished under infection - were strongly associated with mixed lesions in controls (Fig. 5F), suggesting they play roles in containing or regulating combined amyloid–pTau pathology. In fact, with depletion of subset 10 and 14 in cerebral cortex post-infection, we have observed lower abundance of mixed lesions altogether, in favour of pTau (both NeuN^HIGH^ and NeuN^LOW^). Their infection-driven decline could thereby compromise lesion handling or local immune homeostasis. However, the protein aggregate coverage in cerebral cortex remained comparatively stable, as opposed to hippocampus, were total area occupied by these proteinaceous lesions was increased post-infection (Fig. 5D-E).

**Figure 5.**
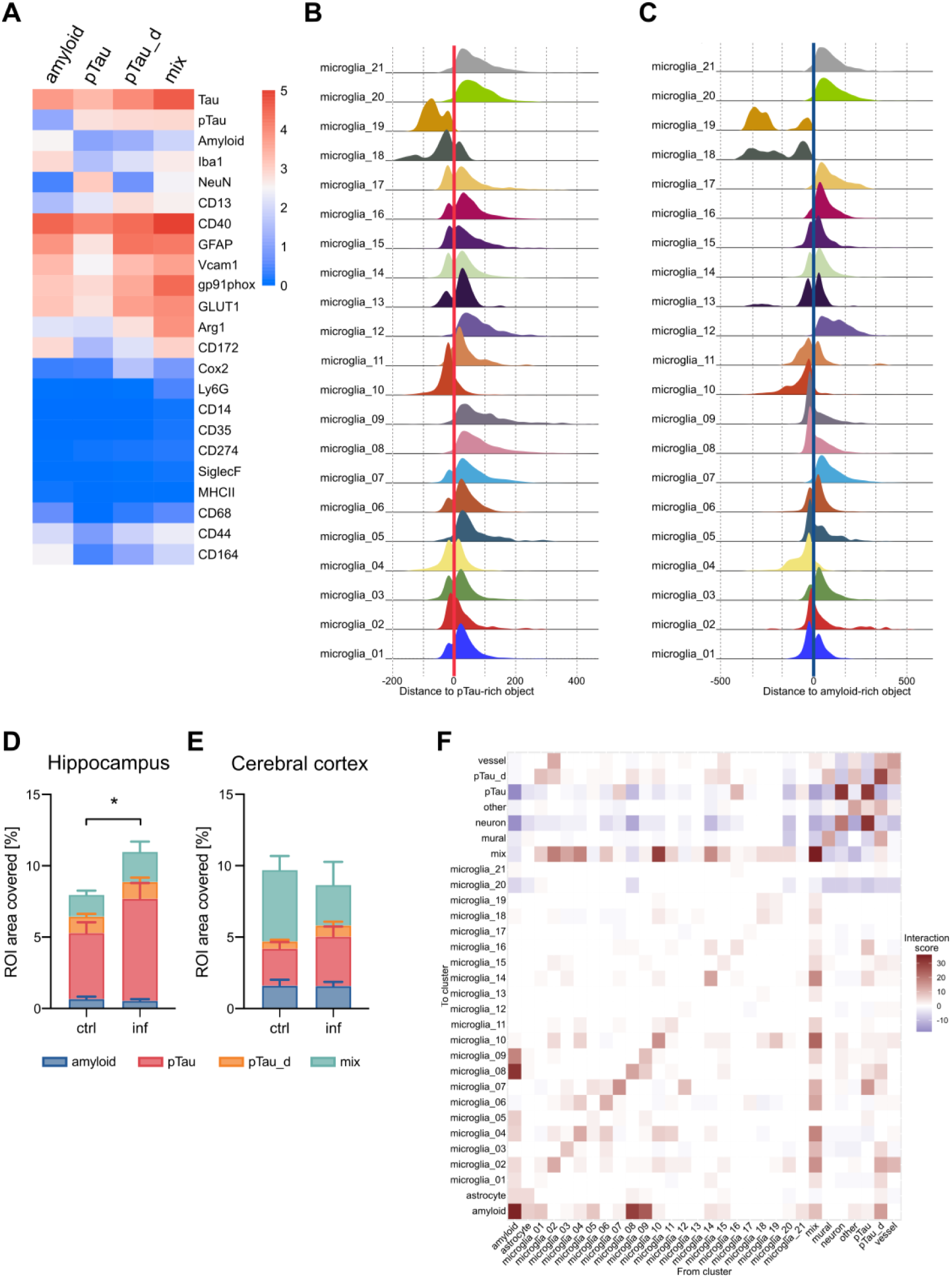
Local cell–aggregate interactions are reshaped in chronically infected hTau-knock-in × 5×FAD mice. **A.** Harmony-integrated UMAP of 60 258 segmented non-cellular objects (see Methods) coloured by four aggregate classes derived by Rphenograph clustering (k = 60): Aβ^HIGH^/pTau^LOW^, pTau^HIGH^/Aβ^LOW^, pTau^HIGH^ + NeuN^HIGH^/Aβ^LOW^, and mixed Aβ^HIGH^ + pTau^HIGH^. **B.** Density plot of microglia (Iba1^HIGH^) subclusters as function of distance to pTau^HIGH^ patches (threshold ≥ 2.35 × background; patch edge = 0 µm). Distance distributions show the number of objects (cells) as a function of their distance from the edge of pTau^HIGH^ patches. **C.** As in b, but for high-amyloid patches (threshold ≥ 1.5 × background). **D.** Percentage of ROI area occupied by pTau or amyloid aggregates in hippocampus and cortex (bars, mean ± s.d.). Cumulative aggregate coverage is higher in infected mice in hippocampus (Mann Whitney two-tailed, p = 0.0357); in cerebral cortex aggregate coverage is unchanged (p = 0.39). **E.** Heat-map of asymmetric interaction scores (imcRtools testInteractions, patch radius = 3 cells, expansion = 20 px) between the 21 microglial subclusters, other major cell classes, and each aggregate class. Positive enrichment (red) and spatial avoidance (blue) are coded by the summed significance value (Σ sigval; Methods). Infected tissue shows strengthened contacts between disease-associated microglia subclusters and mixed Aβ/pTau aggregates, and reduced contacts between NeuN^HIGH^ debris and homeostatic microglia. Aggregate segmentation, patch detection, proximity and interaction analyses were performed as detailed in Methods; low-frequency patches (< 6 px) were excluded.

Interestingly, the NADPHox^HIGH^ subsets 18 and 19, similar to the highly activated “flash-and-burn” microglia *in vitro*, were strongest infiltrators of both pTau and amyloid rich regions, but their interactions scores with these types of aggregates remained modest. In the hippocampus, microglia 20 remained relatively distant from amyloid/pTau^HIGH^ lesions, with microglia subset 1 as the dominant hippocampal responder to the protein aggregates. These differences reinforce our earlier observation on the region-specific differences in microglial heterogeneity and functionality.

### Linking *In Vitro* and *In Vivo* Pathogen Effects

These *in vivo* data validate and extend the *in vitro* findings in a disease-relevant setting. The selective depletion of moderately activated/regulatory subsets (microglia 10, 14) in the cortex under *Pg* infection strongly correlates with the in vitro phenomenon in which *Pg* WT suppresses Arg1^HIGH^, IL-10^HIGH^ microglia. Moreover, the infiltration of more proinflammatory or oxidative subpopulations around amyloid/pTau aggregates parallels the *“flash-and-burn”* phenotypes induced by gingipains *in vitro*. Although hippocampal shifts were subtler, the in vivo data highlight region-specific vulnerabilities—potentially aligning with early cortical AD pathology or distinct local microenvironments. Although *Tf* was not assessed *in vivo*, the partial parallels in hippocampal microglia underscore how different pathogens might preferentially reprogram distinct brain regions if tested in multi-infection models.

Overall, chronic *Pg* infection in an AD-prone environment (5xFAD x hTau KI mice) leads to a net erosion of regulatory microglial subpopulations critical for immunomodulation (Arg1^HIGH^, IL-10^HIGH^, MERTK^HIGH^), while fostering subsets that localize around pathological aggregates, potentially exacerbating lesion growth or neuroinflammation. These findings provide novel region- and lesion-specific insights into how oral pathogens may synergize with AD pathology and strengthen the case for therapeutic gingipain inhibitors or other antimicrobial strategies aimed at preserving beneficial microglial subsets in aging brains.

## DISCUSSION

Microglia, the resident immune cells of the CNS, have increasingly been recognized as pivotal regulators of AD, capable of both protective and detrimental effects depending on their activation states and the local environment. Advances in single-cell profiling - particularly single-cell RNA-sequencing and high-dimensional cytometry - have revealed a complex spectrum of microglial phenotypes, challenging the once-binary view of “resting” *vs.* “activated” microglia^37,38^. Even in homeostatic conditions, microglia are a highly heterogeneous and adaptable population, with their diversity shaped by regional specialization, proximal cell signals, and systemic influences^30,39^. Despite these insights, much remains unknown about how oral pathogens, especially *Pg* and *Tf*, drive microglial diversity at high resolution. Our study addresses this critical gap by systematically dissecting pathogen-specific microglial phenotypes both *in vitro,* with comprehensive CyTOF analyses, and *in vivo*, using imaging mass cytometry in a mouse model recapitulating hallmarks of AD.

We identified 38 distinct microglial clusters in our *in vitro* experimental model using SIM A9 murine cell line, reported to harbour primary-like phenotypes. *Pg* WT induced robust inflammatory bursts, potentially exacerbating AD-like pathology *via* excessive NF-κB activation^40,41^. Further supporting our data, a previous report highlighted *Pg*’s ability to activate PAR2, resulting in preferential up-regulation of proinflammatory mediators, while not significantly elevating regulatory molecules such as IL-10, effectively impairing microglial self-regulatory mechanisms^42–44^. Conversely, *Tf* drove a faster but more sustained dampening consistent with an “exhausted” phenotype - indicating that specific virulence factors (e.g., gingipains, LPS, fimbriae in *Pg*, the S-layer and plethora of proteases in *Tf*) differentially skew microglial function. Comparisons with isogenic mutants (*Pg* KRAB, *Tf* dS) underscored that discrete bacterial proteins act as “molecular levers,” pivoting microglia between proinflammatory or moderately immunoregulatory states. Such high-dimensional mapping - unprecedented for these pathogens - refines a field where prior work relied on incomplete marker sets or generalized infection models, obscuring the finer details of microglial dynamics.

Our data also clarifies that the presence or absence of gingipains (in *Pg*) or the S-layer (in *Tf*) consistently dictates both the severity and trajectory of microglial activation. Our data support the hypothesis that gingipain-mediated microglial activation intersects with established AD pathways - enhancing oxidative stress (NADPH oxidase, iNOS) and upregulating TLR4, MHC II, co-stimulatory molecules (CD86)^45–47^. These disease-associated microglia have been implicated in AD murine models, often converging around plaques and fuelling ongoing neuroinflammation^48,49^. *Pg* strain lacking gingipains (*Pg* KRAB) exhibited blunted inflammatory states, preserving regulatory subsets such as CD73^HIGH^ or IL-10^HIGH^ microglia - crucial for immunoregulation. Meanwhile, *Tf* S-layer deficiency abrogated its capacity to elevate antigen-presenting (CD40, MHC II) or checkpoint-rich (PD-L1/CD274) microglia suggesting that certain checkpoint-oriented phenotypes require intact S-layer signals^50^. While immune checkpoints typically limit T-cell activation, they could paradoxically maintain smouldering inflammation if they curb microglial–T-cell clearance of misfolded proteins^51–54^. Together, these observations reinforce that microglial phenotypes are carefully orchestrated by pathogen proteins, guiding novel therapeutic strategies (e.g., gingipain inhibitors).

*In vivo*, we employed imaging mass cytometry in hTau-knock in 5xFAD mice to see whether the pathogen-driven *in vitro* phenotypes persist under AD-like conditions. Remarkably, chronic *Pg* infection selectively reduced cortical microglial clusters (e.g., subclusters 10, 14) characterized by moderate MHC II, CD86, and IL-10, mirroring our *in vitro* finding that gingipains erode homeostatic or “mixed” microglia, thus impairing regulatory mechanisms and further propelling AD-like pathology. Intriguingly, prior reports indicate that other chronic infections (e.g., COVID-19) also heighten proinflammatory responses at the expense of moderate immunomodulatory phenotypes, underscoring that sustained inflammatory stimuli frequently displace beneficial or intermediate states^55–57^. Furthermore, we observed that these moderately activated subpopulations (high levels of CD14, CD35, CD68, MERTK) interacted strongly with mixed aggregates (amyloid^HIGH^ and pTau^HIGH^), implying they attempt to phagocytose or regulate lesion growth without causing neurotoxicity^58,59^. Their depletion under infection likely accelerates neurodegenerative lesions, echoing the synergy between chronic inflammation and protein aggregation.

Our spatial analyses confirmed region-specific differences in microglial subcluster composition—cortex vs. hippocampus - consistently aligning with prior hypotheses that region-specific microglial makeups underlie differential vulnerability to aging and pathological stressors^60–62^. Since hippocampal changes were comparatively subtle here, we posit that the relatively advanced disease stage (∼11 months) in our 5xFAD x hTau knock-in mice might eclipse potential earlier hippocampal shifts. Future studies employing younger mice or extending the infection timeline could better distinguish AD *vs*. infection-specific microglial alterations in the hippocampus.

These data tighten the link between oral pathogen infection and AD-relevant microglial reprogramming, bridging two previously disconnected research domains: periodontal infection and microglial heterogeneity in AD. Therapeutically, our findings resonate with the promise of gingipain inhibitors and highlight potential benefits of targeting *Tf*–specific factors (S-layer), especially if multi-pathogen synergy ultimately proves critical in AD progression. Future work exploring co-infections (e.g., *T. denticola*, *F. nucleatum*, or fungal/viral species) may uncover further microglial states or synergies that accelerate - or possibly mitigate - AD pathogenesis.

Our findings also highlight the power of spatial proteomics—unlike conventional immunofluorescence, this approach enabled precise mapping of microglial subtypes potentially critical to AD pathogenesis. By capturing multiple functional markers simultaneously, our mass cytometry pipelines exceed single-marker approaches, unveiling unprecedented microglial plasticity under infection stress. This broad, integrated view illuminates how discrete microglial subsets fluctuate depending on which bacterial virulence factors predominate - and how these shifts coincide with AD lesion development. Additionally, our protein-level focus circumvents transcriptome-proteome mismatches that can complicate single-cell RNA-seq analyses in aged or diseased brains^63–65^. Hence, this study provides a conceptual and methodological platform for dissecting oral pathogen– microglia interactions, bolstering the rationale that targeting microbial virulence (gingipains, S-layer) might help alleviate microglial dysregulation and decelerate AD progression.

Nonetheless, open questions remain. Longitudinal *in vivo* imaging or serial sampling will clarify whether pathogen-induced microglial changes directly accelerate plaque or tangle formation, and whether early blockade of gingipains or S-layer can spare crucial immunoregulatory microglia and improve cognitive outcomes. Overall, our findings accentuate that microglia, far from bystanders, actively detect bacterial virulence signals in their vicinity and transition into states consistent with heightened or chronic neuroinflammation. By reconstructing these pathogen-specific microglial responses at single-cell resolution—both *in vitro* and in an AD-like environment - we illustrate how oral pathogens globally alter microglial heterogeneity, merging periodontal research with the evolving landscape of AD immunopathology. This synergy reveals actionable mechanistic pathways for future interventions aimed at neutralizing virulence factor–driven neuroinflammatory cascades, offering rational therapeutic designs to complement established anti-amyloid or anti-tau strategies.

Finally, by coupling a high-dimensional *in vitro* model with a 5xFAD x hTau KI system, we demonstrate that *Pg* and *Tf* exploit distinct virulence factors - gingipains and S-layer - to reprogram microglial subsets along proinflammatory, antigen-presenting, or immunoregulatory axes, validating that infection-driven heterogeneity indeed applies under AD-like conditions. These insights move beyond monolithic views of “activated” microglia, identifying pathogen-derived molecular “switches” as promising targets to contain infection-driven neuroinflammation in Alzheimer’s disease.

## Materials and Methods

### Bacteria culture

*Porphyromonas gingivalis (Pg)* W83, gingipain-deficient *Pg* ΔKgpΔRgpAΔRgpB in W83 background (KRAB), *Tannerella forsythia (Tf)* ATCC 43037 and non-S-layer-producing *Tf* were cultured on BD BBL Brucella agar (Beckton Dickinson; 211086) plates supplemented with hemin (SigmaAldrich, H9039), vitamin K (SigmaAldrich, M5625), and 5% defibrinated sheep’s blood, under anaerobic conditions at 37°C. For *Tf*, agar medium was additionally supplemented with N-acetylmuramic acid (0.01 mg/ml; SigmaAldrich, A3007). Bacteria were passaged onto fresh agar every 48h (*Pg*) or 120h (*Tf*). For the *Pg* KRAB strain, agar plates contained tetracycline (1 µg/ml; SigmaAldrich, T7660) and erythromycin (5 µg/ml; SigmaAldrich, E5389).

Bacterial suspensions for microglia infections were prepared by suspending freshly cultured bacteria (≤48h for *Pg*, ≤72h for *Tf*) in sterile PBS, followed by centrifugation (4600 rpm, 10 min, 4°C). After resuspension in PBS, bacterial aggregates were removed by low-speed centrifugation (800 rpm, 5 min, 4°C). Optical density (OD_600_) was measured, and colony-forming units (CFUs) were estimated (*Pg*OD600 = 1 corresponding to 1 × 10^9^ CFU/ml; *Tf* OD_600_ = 1 corresponding to 6.8 × 10^8^ CFU/ml).

### Cell line culture

The murine microglial SIM-A9 cell line (ABM, T0247) was cultured in DMEM/F12 (ThermoFisher, 31331093) containing 10% FBS (ThermoFisher, A5256801), 5% horse serum (ThermoFisher, 16050), 100 U/ml penicillin, and 100 µg/ml streptomycin (ThermoFisher, 15140122), at 37°C in a 5% CO₂ humidified incubator. Cells were passaged by pooling suspension and adherent fractions, followed by trypsinization, centrifugation (280g, 10 min, RT), and resuspension in fresh medium. Cells were seeded at a 1:10 dilution; passages 5 and 6 were used for infection experiments. Cell density was measured using trypan blue exclusion on a Bürker chamber.

### Microglia infection

SIM-A9 cells were seeded at 5.33 × 10⁵ cells/ml in 6-well plates and allowed to adhere overnight (16h). Before bacterial challenge, the medium was replaced with serum- and antibiotic-free DMEM/F12, and cells were incubated for an additional 2h. Bacteria were introduced at multiplicities of infection (MOI) of 5 or 10. Cells collected at 0h (pre-infection) were washed with PBS, while infected cells were harvested at 24h and 48h.

Cell suspensions were centrifuged at 300g for 5 min in RT and resuspended in PBS. Samples were centrifuged (300g, 5 min in RT), resuspended in Maxpar Cell Staining Buffer (Standard Biotools, 201068). Fixing and barcoding were perfomed according to the manufacturer’s instruction. Cells were fixed in 4% paraformaldehyde (PFA; ThermoFisher, 28908) diluted in PBS for 5 min, washed in Barcode Perm Buffer (Standard Biotools, 201057) and stained with barcodes: Pd-102, Pd-104, Pd-105, Pd-106, Pd-108, Pd-110 (Cell-ID 20-plex Pd Barcoding kit, Standard Biotools, 201060) in Barcode Perm Buffer for 30 min in RT. Some samples were additionally stained with 20 nM cisplatin (cisPt195, Neonest AB) to increase sample pooling capacity. Barcoded cells were then washed in Maxpar Cell Staining Buffer and pooled together for antibody staining.

### Polymer conjugation

All antibodies, except those purchased directly from Standard Biotools, were conjugated to metal-chelating polymers in-house. Conjugating kits for: 144Nd, 145Nd, 146Nd, 147Sm, 148Nd, 149Sm, 150Nd, 152Sm, 155Gd, 158Gd, 160Gd, 161Dy, 162Dy, 163Dy, 165Ho, 166Er, 167Er, 168Er, 169Tm, 173Yb, 175Lu, 176Yb were purchased from StandardBiotools, while kits for: 115In, 170Er and 171Yb were purchased from Ionpath. All conjugations were performed according to the respective manufacturer’s instruction. For each conjugation, antibodies (100 µg per conjugation) were used without bovine serum albumin (BSA). Antibody protein concentrations were measured using NanoDrop spectrophotometer at 280 nm before and after conjugation. Final concentrations were adjusted to 0.5 mg/ml in Antibody Stabilizer with HRP-Protector™ (Candor Bioscience, 220 050) for suspension staining, or in manufacturer-provided buffer (Ionpath) for imaging mass cytometry. All conjugates were stored at 4°C.

### Antibody panels: titration and validation

Prior to experimental use, all antibodies were rigorously titrated on SIM A9 cells. Cells were fixed, then blocked using anti-mouse CD16 antibody (1 µg per 3 million cells, BioLegend, cat. 101302) for 10 min at room temperature (RT). Serial antibody dilutions ranging from 1:100 to 1:3200 were applied to the cells and incubated for 30 min at RT. Cells were subsequently washed with Maxpar Cell Staining Buffer and fixed again with 4% PFA for 5 min at RT. For antibodies targeting intracellular proteins (TNFα, IL-10, IL-12p40, TGFβ, Arg1, NADPH oxidase, and iNOS) cells underwent permeabilization with Maxpar Perm-S Buffer (Standard Biotools, 201066) for 30 min at RT, followed by intracellular antibody staining for 30 min at RT. After staining, cells were washed once more in Maxpar Cell Staining Buffer, fixed in 1.6% PFA for 10 min, and nuclei were stained with Cell-ID Interacalator (Standard Biotools, 201192A) at 31.25 nM for 1h. For signal acquisition, cells were suspended in Maxpar Cell Acquisition Solution Plus (Standard Biotools, 201244). For antibody selectivity confirmation, SIM A9 cells were stimulated with 100 ng/ml recombinant murine IFNy (StemCell, 78021.1) for 16h prior to antibody staining. Data was collected as described in the Data Collection section. The optimal dilution for each antibody was identified as the dilution providing clear signal intensity above the minimal detectable threshold but significantly below the maximal point of the dynamic range (Supplementary Fig. 1A-B).

For imaging mass cytometry, staining was performed as described below using OCT-embedded brain sections (thickness 10 µm) prepared from C57BL/6 wild-type mice. For titration procedure, 1:50 – 1:200 dilutions were tested, and all antibodies were incubated with tissues for 16h in 4°C. When antibody signal-to-noise ratio was deemed sufficient on visual inspection, antibody selectivity was inspected based on previously established expression patterns and tissue architecture. Titrated antibodies that passed this initial inspection were then validated on liver sections (thickness: 10 µm) selected for their abundance of Kupfer cells. Antibodies that exhibited differential and anatomically appropriate expression patterns across brain and liver tissues were confirmed as validated and included in subsequent imaging mass cytometry analyses (Supplementary Fig. 1D).

### Data collection

Barcoded and pooled SIM A9 were blocked with anti-mouse CD16 (1 µg/3 million cells, BioLegend, cat. 101302), and stained with antibody cocktail (Supplementary Table 1, surface markers). Cells were washed briefly in Maxpar Cell Staining Buffer, fixed again (4% PFA in PBS, 5 min in RT), permeabilised with Maxpar Perm-S Buffer and then stained with intracellular marker antibody cocktail (Supplementary Table 1) diluted in Maxpar Perm-S Buffer for 30 min in RT. Then, cells were washed once again in Maxpar Cell Staining Buffer, fixed in 1.6% PFA for 10 min and nuclei were stained with Cell-ID Interacalator at 31.25 nM for 1h.

For signal acquisition, cells were suspended in Maxpar Cell Acquisition Solution Plus and strained (Falcon, 352235). Cells were pelleted (800 g for 2 min in 4°C) and pellets were loaded onto CyTOF instrument, where they were suspended in Maxpar Cell Acquisition Solution Plus at 0.75 mln/ml density. EQ^TM^ Six Element Calibration Beads (33,000 beads/ml; StandardBiotools) were added and signal was recorded at speed of 200 – 350 events/s using CyTOF XT (Standard Biotools), operated by CyTOF software 8.1.0 (Standard Biotools). Bead-based signal normalisation and initial event debarcoding was performed using CyTOF software and Debarcoder algorithm with provided barcode key as a .csv file with Maximum Amhalanobis Distance = 10. Generated data was then expored as .fcs files for further processing.

### Data preprocessing

Initial processing was performed in FlowJo v10.8.1 (Beckton Dickinson). Samples were gated for cisplatin-based barcodes, and EQ bead events (140Ce) were excluded. Doublets (Ir191 vs Ir193) and debris (Ir191 vs event length) were also removed (Supplementary Figure 1C). Clean .fcs files were imported into R (SingleCellExperiment v1.26.0, flowCore v2.16.0), transformed using an inverse hyperbolic sine function (asinh, co-factor = 5). Subsequently, samples were downsampled to equalize event counts across biological replicates, resulting in a final dataset comprising 745,000 cells in total. No spillover compensation was performed on this dataset.

### Clustering

Uniform Manifold Approximation and Projection for Dimension Reduction (UMAP) was generated. Phenoannoy^66^ (0.1.0, at k=800) was used for unsupervised clustering, as number of events exceeded computational power required to employ Rphenograph package, while FLOWsom package did not provide sufficient resolution to highlight microglial heterogeneity *in vitro*.

### Animals

Generation of 5xFAD x hTau KI mouse model was described in Barendrecht et al 2023^36^. All animal experiments were approved by the responsible animal ethics committee of the state of Saxony-Anhalt, Germany (Landesverwaltungsamt Sachsen-Anhalt, Department of Consumer Protection and Veterinary Affairs, Halle (Saale), Saxony-Anhalt, Germany) under the following approval numbers: 42502-2-1371 MLU and 42502-2-1670 IZI.

Fifty µl containing either 10^9^ CFU *Pg* (strain ATCC 33277) in 2% methylcellulose (experimental group) or 2% methylcellulose in PBS (control group) was administered into the oral cavity three times per week for 22 weeks, starting at the age of six months. At the age of 12 months, mice were sacrificed by CO_2_ inhalation followed by blood withdrawal and were then transcardially perfused with PBS (pH 7.4). Sample collection and preparation for immunofluorescence staining was carried out according to Barendrecht et al 2023.

### Immunofluorescent staining

The staining was performed on free-floating 30 µm sagittal brain sections. Sections were incubated with TBS, 0.3% Triton-X-100, 5% normal goat serum for 30 min followed by the incubation with Iba-1 (1:250; FUJIFILM Wako Pure Chemical Corporation, Japan) at 4°C overnight in a humid chamber. The following day sections were washed with TBS and incubated with Cy3-labelled goat anti-rabbit secondary antibody (1:250, Dianova) for 60 min. After washing with TBS, the sections were placed under the microscope on slides coated with protein glycerol, air-dried and mounted in mounting media.

### Fluorescence microscopy

Stained brain tissue sections were examined with an AxioScan.Z1 microscope (Zeiss, Göttingen, Germany) equipped with a Colibri 7 light source. Excitation / Emission filters for Cy3 were at 555 and 595 nm, respectively. Pictures were taken by means of Axiocam 506 and a Plan-Apochromat objective (20×/0.8) and exposure time of 20 ms for Cy3. Images were digitized by means of ZEN blue 3.4 software. Brain regions of interest (cortex and hippocampus) were extracted before microglia analysis by using this software.

### Microglia Analysis

All immunofluorescence images were processed by a developed macro for ImageJ/Fiji^67,68^. In a preprocessing step, the Cy3 channel was normalized and enhanced in contrast as well as cleaned of artifacts from the stitching procedure of the slide scanner or uneven illumination. The Cy3 channel was binarized with a local mean thresholding algorithm using the same set of parameters for every single image. In the last step, information from the channels is being extracted regarding the object’s sizes and the count of objects for each image with the ImageJ/Fiji tool *Analyze Particles*.

### Tissue staining

Tissue slides were thawed by first incubating at −20°C for 2 hours, followed by 20 min at 4°C. OCT compound was removed by brief incubation in PBS at 4°C. Subsequently, antigen retrieval was performed using EnVision FLEX Target Retrieval Solution High pH (Agilent Dako, K8004; pH = 9), incubated for 15 min at 96°C. Slides were then cooled down to 65°C, briefly rinsed in PBS at room temperature (RT), and permeabilized by incubation in PBS containing 0.5% Triton X-100 for 20 min at RT. Following a PBS wash, tissues were blocked in 4% BSA in PBS for 1 hour at RT.

After blocking, samples were stained with a freshly prepared antibody cocktail (Supplementary Table 2), diluted in 0.5% BSA in PBS, and incubated for 20 hours at 4°C. Unbound antibodies were removed through two sequential 8-minute washes in PBS containing 0.5% Triton X-100 with gentle agitation. Finally, nuclei were stained for 20 min in RT with Cell-ID Intercalator at 156 nM in PBS. Slides were then washed twice in Mili-Q water and air-dried.

### Scanning

Tissues were scanned using Hyperion Imaging System (Standard Biotools) coupled with a Helios mass spectrometer (Standard Biotools), CyTOF software (version 7.1.16401.0). Complete hippocampus area (1.19 mm^2^) and two regions of interest (ROIs, area 1.19 mm^2^ each) from the cerebral cortex dorsal to hippocampus were scanned. Tissue ablation was performed at 1 dB power with a frequency of 200 Hz and an X–Y step size of 1 px (1 px = 1 µm), corresponding to the maximum resolution of the instrument. Signal intensities from each channel were recorded for every scanned pixel and exported as .txt files for further processing.

### Spillover correction and denoising

A compensation matrix was generated using an in-house antibody microarray. Individual air-dried antibody droplets were scanned, and background signals were measured across all other panel channels, as previously described^69^. Raw intensity data were imported and asinh-transformed with a cofactor of 5. Pixel-level debarcoding was performed using the CATALYST (v1.28.0) package. Misidentified pixels were removed, and spillover was quantified using a channel-wise compensation matrix (Supplementary Fig. 2A).

.txt files were processed using the steinbock pipeline. Files were converted to .tiff format with hot pixel filtering set to 50. Images were subsequently loaded using imcRtools (v1.10.0) and cytomapper (v1.16.0). Signal spillover was corrected across all channels except Ir191 and Ir193, using the previously generated matrix. Both pre- and post-compensation images were visually inspected for artifacts and saved as 32-bit .tiff files.

Denoising and background removal were performed using the self-supervised IMC-Denoise package^70^. IMC-Denoise environment was created (python 3.6, tensorflow 2.2.0, keras 2.3.1). A randomized subset of 20 ROIs was used for training. Model performance was visually assessed in channels with signal loss exceeding 0.02 normalized intensity. Channels CD274, SiglecF, and MHCII were excluded from denoising due to insufficient signal-to-noise contrast.

### Object segmentation

Cell segmentation was performed using a hybrid Ilastik–CellProfiler workflow^71–73^. Random 250 × 250 px patches were generated from spillover-corrected and denoised images for pixel classifier training in Ilastik (v1.4.0). Three probability channels were defined using visually validated thresholds: channel 1 (nuclei) - trained with Ir193/Ir195 channels in training patches (signal threshold: 37), channel 2 (cytoplasm) - trained with NeuN (signal threshold: 6), GFAP (signal threshold: 23), Iba1 (signal threshold: 9), S100B^HIGH^ (signal threshold: 23) and GLUT1^HIGH^ (signal threshold: 21); and channel 3 as background channel. The resulting probability maps (Supplementary Fig. 2B) were then imported into CellProfiler (v4.2.7) using custom pipelines derived from the steinbock package. The nuclei channel was used as a seed (object size: 3–25 px) for cytoplasmic mask propagation (Supplementary Fig. 2B). To preserve the morphological features of CNS cells, the “Fill holes in identified objects” function was disabled during segmentation.

Protein aggregates segmentation was performed in a smiliar manner. Ilastik was used to generate probability maps: channel 1 was pTau^HIGH^ regions, channel 2 - amyloid^HIGH^, and channel 3 as background. Subsequently, these probability maps were imported into CellProfiler, where the combined signal from Channels 1 and 2 was utilized for precise identification and segmentation of protein aggregates.

### Data cleanup

The processing workflow was adapted from Windhager et al. with modifications^74^.

Properties of objects identified as described above were evaluated using the tools in the steinbock package (0.16.1). In short, object intensities and sizes were measured and exported to csv file. Data (tiff images, segmentation masks and single-cell data) were read in Rstudio environment (R 4.4.3, Rstudio 2024.12.1 Build 563). Signal was asinh-transformed (co-factor =1), and signal-to-noise ratios were quality checked. Objects with areas smaller than 4 px or larger than 2500 px (cell-like segmentation) or larger than 3000px (aggregates) were excluded from downstream analyses. UMAP dimensionality reduction was performed using scater (v1.32.1) on all markers except DNA1/2. Batch correction was conducted using Seurat v.5.2.0 and SeuratObject v.5.0.2 packages^75^. Data were split by slide ID, integrated via reciprocal PCA (k = 20), and reduced to principal components for batch-corrected UMAP generation. Batch-corrected integration quality was visually assessed (Supplementary Fig. 2C).

Aggregate segmentation data were batch-corrected using harmony v.1.2.3^76^, which exhibited improved biological fidelity compared to Seurat in this context (Supplementary Fig. 5). PCA computations were supported by BiocSingular.

### Clustering

Cell and protein aggregate typing were performed through unsupervised clustering using the Rphenograph package (v.0.99.1.9003)^66^. The initial clusters from the cell dataset were annotated at k=50 for n = 78 622 events. The resulting initial clusters were sorted into the following cell types (hence called metaclusters) based on known patterns of marker expressions: neurons (NeuN^HIGH^), blood vessels (GLUT1^HIGH^), astrocytes (GFAP^HIGH^), microglia (Iba1^HIGH^), mural cells (CD13/Cox2^HIGH^) or other cell types that did not distinctly match any of these predefined categories. Due to imaging resolution constraints (minimum scan resolution of 1 µm), some degree of antigen signal overlap or cross-contamination (i.e. GFAP^+^ in Iba1^HIGH^ cell types) was considered acceptable and did not preclude accurate annotation. Subsequently, microglia-like objects from the broader dataset were isolated and further subclustered using Rphenograph with k = 60, resulting in the identification of 21 distinct microglial subclusters.

Aggregate typing was performed similarly using Rphenograph (k = 60, n = 60 258). Aggregates were then systematically assigned one of four categorical labels based on specific marker positivity profiles: amyloid (amyloid^+^, but pTau^-^ objects), pTau (pTau^+^, NeuN^-^), pTau_d (pTau^+^ and NeuN^+^) or mixed aggregate (amyloid^+^ and pTau^+^).

### Spatial interactions

Cell-to-cell interactions were evaluated by constructing spatial graphs using imcRtools’ buildSpatialGraph function, with an expansion threshold of 40 px. For each object, neighboring cells were identified and annotated according to their metacluster classification, which enabled the organization of cells into six distinct regional microenvironments. The compositional structure of these regions—both at the level of broad metaclusters and finer microglial subtypes - was visualized using the lisaClust package^77^.

Next, region-to-region spatial contexts (SCs) were defined using a k-nearest neighbor (k-NN) approach (k = 40). To ensure representation across multiple samples, only SCs with ≥200 total cells and identified in ≥5 ROIs were retained.

To quantify interactions between defined clusters—including interactions between cells and protein aggregates - proximity-based analysis was performed using the testInteractions function within imcRtools package, based on proximity expansions (k = 40) in 3-cell-radius patches. Resulting interaction scores were color-coded according to the summed significance values (sigval), with positive enrichment interactions visualized in red and spatial avoidance represented in blue.

### Aggregate proximity analysis

To characterize the cellular composition surrounding pathological protein aggregates, datasets were merged following cell and aggregate clustering. High-density amyloid and pTau regions were identified using the patchDetection function in *imcRtools*, with detection thresholds set at ≥1.5× the relative signal intensity for amyloid and ≥2.35× for pTau. A minimum patch size of 6 pixels was applied, followed by a 2-pixel expansion to encompass neighboring structures. Patch boundaries were validated by comparing patch area distributions against the corresponding segmented object sizes, ensuring consistency in detection criteria.

To assess spatial proximity between cells and pathological regions, distances from each segmented cell object to the nearest aggregate patch were calculated using the plotSpatial function. Distance distributions were then stratified by cell type, enabling comparative analyses of microglial, astrocytic, and neuronal positioning relative to amyloid or pTau pathology.

### Statistical analysis and data visualisation

Cluster abundance distributions were first assessed for normality using the Shapiro–Wilk test. As most datasets did not meet the assumption of normality, non-parametric Kruskal–Wallis tests were applied for group comparisons, followed by Dunn’s post hoc correction. P-values were adjusted for multiple testing (Padj). Statistical analyses were performed using stats v4.4.3, FSA v0.9.6 with dplyr v1.1.4 for necessary data transformations.

For data visualisation, GraphPad Prism v.10 or the following R packages were employed: cowplot v1.1.3, dittoSeq v1.16.0, ggplot2 v3.5.1, ggridges v0.5.6, pheatmap v1.0.12. Parallel computations were performed with BiocParallel v1.38.0. All R packages were downloaded and installed through Bioconductor project (BiocManager v1.30.25).

### Data availability statement

Cytof and IMC datasets have been deposited at doi.org/10.57903/UJ/MQOATZ.

### Code availability statement

All R scripts have been deposited at github.com/Dorlana/MicrogliaRush2025.

## Acknowledgements

The authors thank Prof. Steffen Rossner and Dr. Corinna Höfling from Paul Flechsig Institute for Brain Research, University of Leipzig, Germany for help with scanning fluorescent brain sections. Mass cytometry was performed at the Flow Cytometry Core Facility, University of Bergen.

---

**Extended Data Figure 1.**
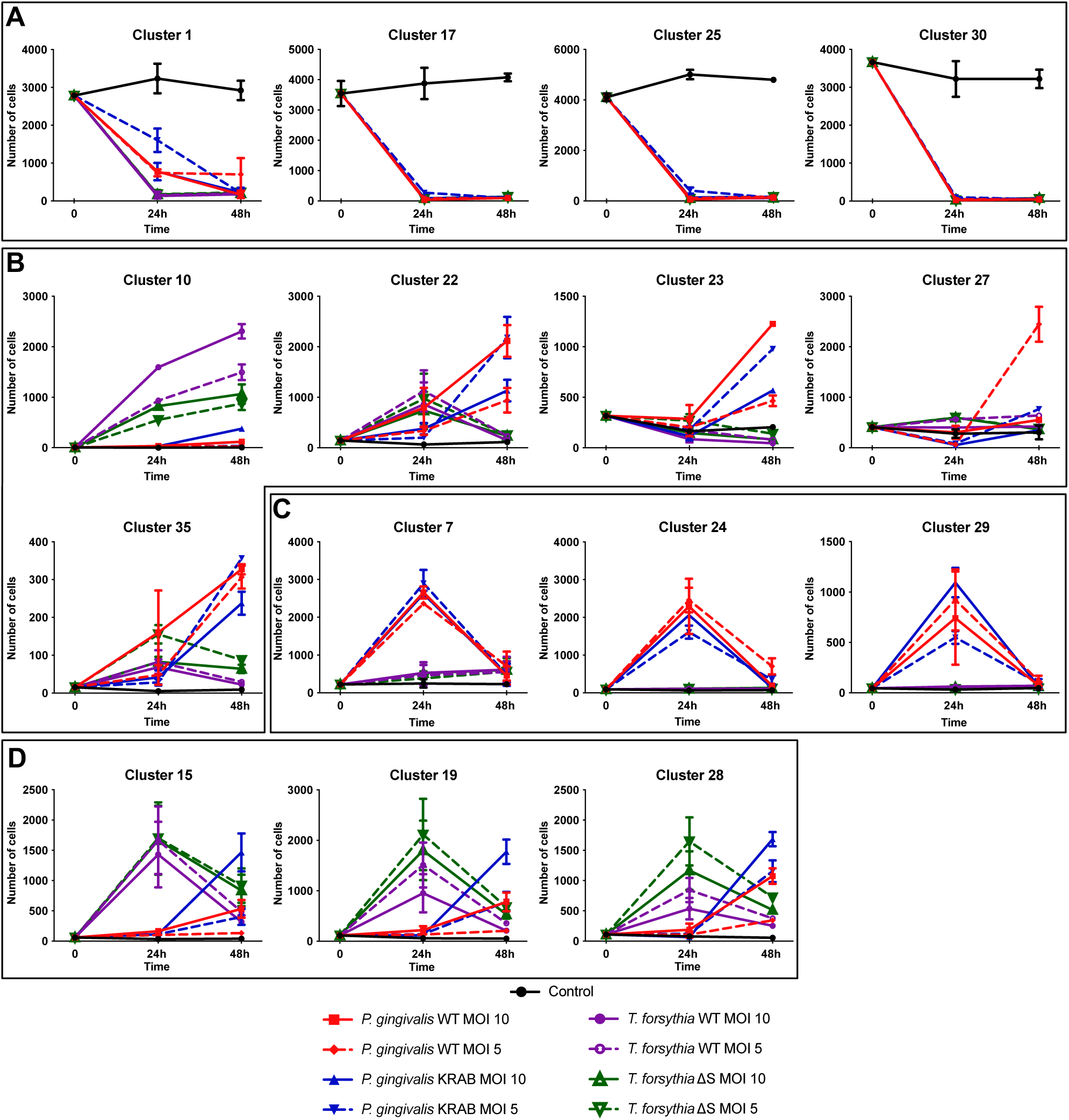
Temporal dynamics of SIM-A9 microglial clusters following infection with P. gingivalis or T. forsythia. Line plots show relative abundance of selected clusters across conditions at 0 h (untreated), and 24 h and 48 h post-infection, following downsampling to equal cell numbers per condition. Each point represents mean of biological replicates ± SEM; lines connect mean values. A. Clusters 1, 17, 25 and 30. B. Clusters 10, 22, 23, 27 and 35. C. Clusters 7, 24 and 29. D. Clusters 15, 19 and 28. E. Clusters 26 and 31. Color-coded legend indicates treatment conditions and pathogen strains (MOI 5 or 10), as in Fig. 2. Cluster-specific responses show distinct temporal trajectories and pathogen sensitivities. Statistical analyses are shown in Fig. 2b and Methods.

**Extended Data Figure 2.**
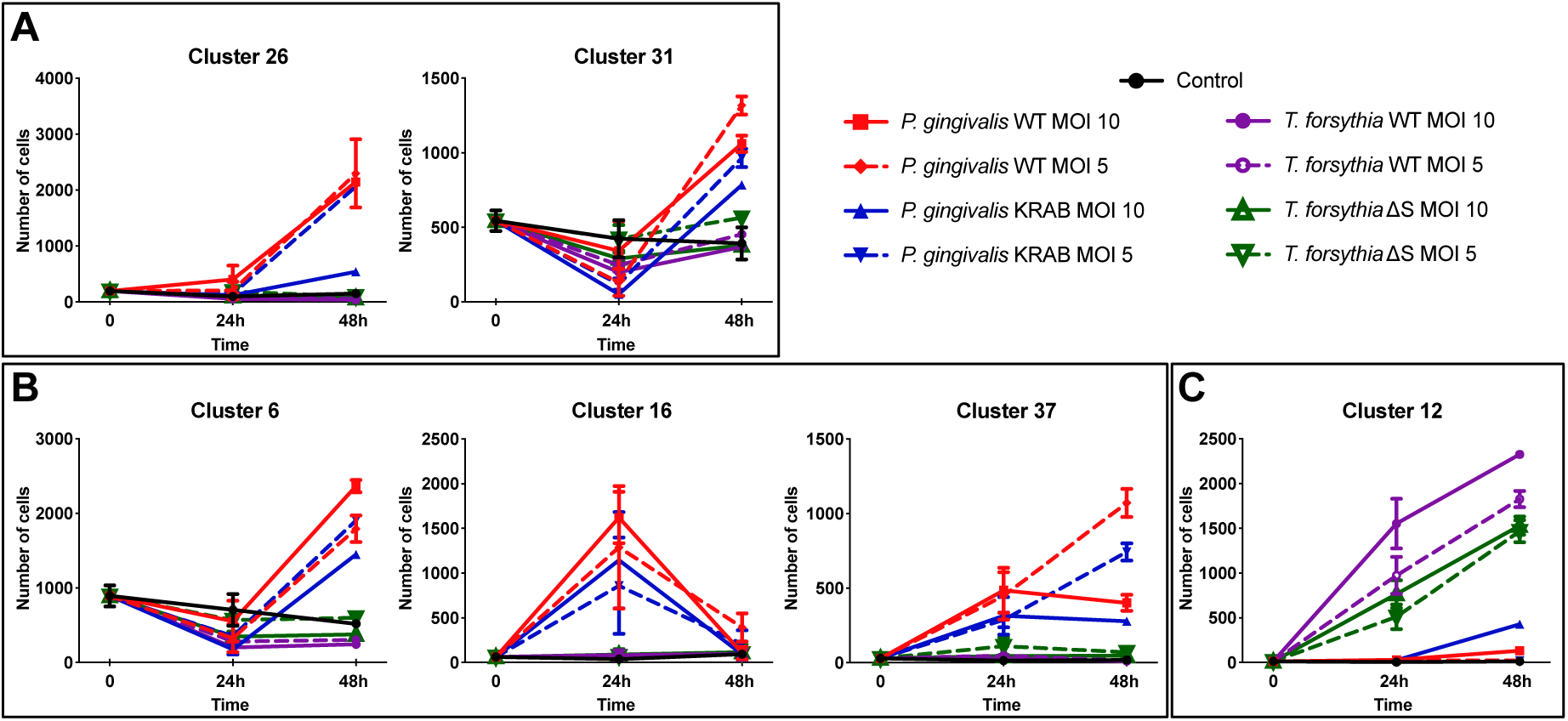
Temporal dynamics of SIM-A9 microglial clusters following infection with P. gingivalis or T. forsythia. Line plots show relative abundance of selected clusters across conditions at 0 h (untreated), and 24 h and 48 h post-infection, following downsampling to equal cell numbers per condition. Each point represents mean of biological replicates ± SEM; lines connect mean values. A. Clusters 26 and 31. B. Clusters 6, 16 and 37. C. Cluster 12. Color-coded legend indicates treatment conditions and pathogen strains (MOI 5 or 10), as in Fig. 2. Cluster-specific responses show distinct temporal trajectories and pathogen sensitivities. Statistical analyses are shown in Fig. 2b and Methods.

**Extended Data Figure 3.**
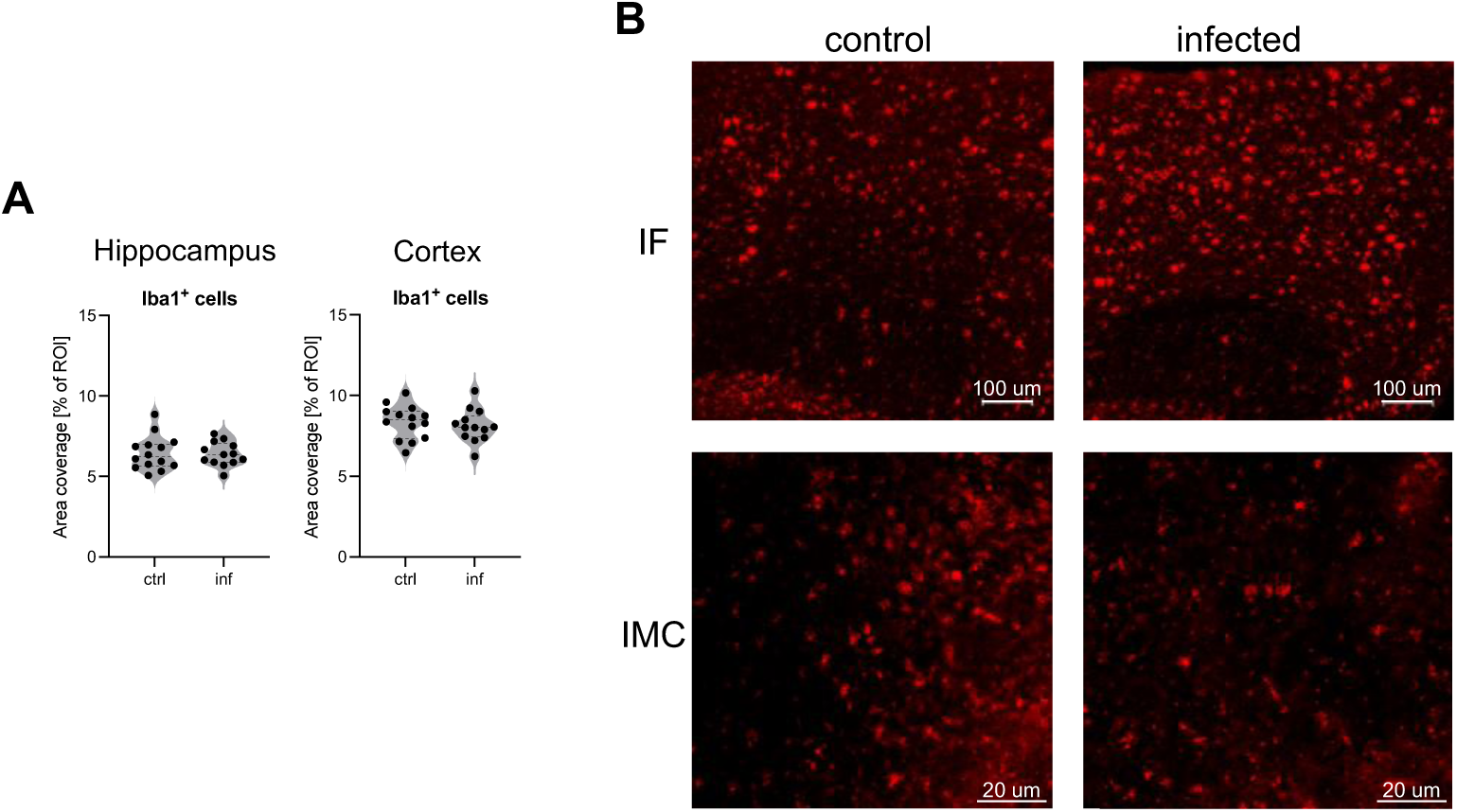
Microglial abundance is unaltered by chronic infection: concordant results from immunofluorescence and imaging mass cytometry. A. Quantification of Iba1⁺ area in the hippocampus (left) and cerebral cortex (right) shows no significant difference between control and P. gingivalis–infected hTau × 5×FAD mice, as assessed by immunofluorescence. Data represent percentage of Iba1⁺ area per brain region. B. Representative images of Iba1⁺ microglia from both control and infected animals, obtained using immunofluorescence (top; scale bar = 100 µm) and imaging mass cytometry (bottom; scale bar = 20 µm), demonstrate comparable Iba1 signal distribution and morphology across methods. Immunofluorescence staining was performed on free-floating 30 µm sagittal brain sections using anti-Iba1 (1:250, FUJIFILM Wako) and Cy3-conjugated secondary antibody (1:250, Dianova). Sections were mounted in glycerol-based media and imaged using an AxioScan.Z1 microscope (Zeiss) with Cy3 excitation/emission (555/595 nm), a 20×/0.8 Plan-Apochromat objective, and 20 ms exposure time. Image processing was performed using ZEN blue 3.4 software. Microglial area analysis was conducted using a custom ImageJ/Fiji macro. Cy3 channel intensity was normalized and contrast-enhanced, followed by artifact correction and binarization using a local mean thresholding algorithm. Iba1⁺ area was quantified using the Analyze Particles tool under uniform parameters across all samples. Both immunofluorescence and imaging mass cytometry approaches yielded consistent results, showing no significant differences in regional microglial abundance between experimental groups.

