## Supplementary materials for "Infection-Specific Reprogramming of Microglia Reveals Distinct Virulence Pathways Linking Periodontal Pathogens to Alzheimer’s Disease"

### Supplementary Data

Supplementary Tables 1 – 2

Supplementary Figures 1 – 5

**Supplementary Table 1.** Antibodies used in the suspension panel (CyTOF, SIM A9 experiment).

| Antigen name | Source | Catalog number | Type | Metal tag | Dilution |
|---|---|---|---|---|---|
| CD45 | Standard Biotools | 3089005B | Monoclone (30-F11) | 89Y | 1/400 |
| TNFalpha | Standard Biotools | 3141013B | Monoclone (MP6-XT22) | 141Pr | 1/100 |
| CD11c | Standard Biotools | 3142003B | Monoclone (N418) | 142Nd | 1/400 |
| CD68 | Biolegend | 137002 | Monoclone (FA-11) | 143Nd | 1/400 |
| CD54 (ICAM-1) | Biolegend | 116102 | Monoclone (YN1/1.7.4) | 144Nd | 1/3200 |
| CD301 | Biolegend | 145702 | Monoclone (LOM-14) | 145Nd | 1/100 |
| IL10 | Biolegend | 505029 | Monoclone (JES5-16E3) | 146Nd | 1/200 |
| TLR4 | Biolegend | 145402 | Monoclone (SA15-21) | 147Sm | 1/100 |
| Iba1 | abcam | ab221790 | Monoclone (epr16589) | 148Nd | 1/200 |
| IL12p40 | BD Biosciences | 554477 | Monoclone (C15.6) | 149Sm | 1/200 |
| CD24 | Standard Biotools | 3150009B | Monoclone (M1/69) | 150Nd | 1/200 |
| CD64 | Standard Biotools | 3151012B | Monoclone (X54-5/7.1) | 151Eu | 1/100 |
| NADPH Oxidase | BD Biosciences | 611415 | Monoclone (53/gp91[phox]) | 152Sm | 1/200 |
| CD274 | Standard Biotools | 3153016C | Monoclone (10F.9G2) | 153Eu | 1/200 |
| CD11b | Standard Biotools | 3154006B | Monoclone (M1/70) | 154Sm | 1/3200 |
| CD40 | Biolegend | 102902 | Monoclone (HM40-3) | 155Gd | 1/200 |
| CD14 | Standard Biotools | 3156009B | Monoclone (Sa14-2) | 156Gd | 1/800 |
| TGF-beta | BD Biosciences | 555052 | Monoclone (A75-2) | 158Gd | 1/200 |
| F4/80 | Standard Biotools | 3159009B | Monoclone (BM8) | 159Tb | 1/400 |
| CD35 | BD Bioscences | 558768 | Monoclone (8C12) | 160Gd | 1/200 |
| iNOS | Standard Biotools | 3161011B | Monoclone (CXNFT) | 161Dy | 1/100 |
| CD44 | Biolegend | 103002 | Monoclone (IM7) | 162Dy | 1/3200 |
| CD172a | BD Biosciences | 552371 | Monoclone (P84) | 163Dy | 1/3200 |
| CX3CR1 | Standard Biotools | 3164023B | Monoclone (SA011F11) | 164Dy | 1/100 |
| MERTK | Biolegend | 151502 | Monoclone (2B10C42) | 165Ho | 1/100 |
| Arginase-1 | BD Biosciences | 610708 | Monoclone (Clone 19) | 166Er | 1/1600 |
| CD124 | BD Bioscences | 551853 | Monoclone (mIL4R-M1) | 167Er | 1/100 |
| CD206 | Biolegend | 141702 | Monoclone (C068C2) | 168Er | 1/100 |
| CD51 | Biolegend | 104102 | Monoclone (RMV-7) | 169Tm | 1/100 |
| SiglecF | BD Bioscences | 552125 | Monoclone (E50-2440) | 170Er | 1/100 |
| CD38 | Standard Biotools | 3171007B | Monoclone (90) | 171Yb | 1/100 |
| CD86 | Standard Biotools | 3172016B | Monoclone (GL1) | 172Yb | 1/800 |
| CD200R | Biolegend | 123902 | Monoclone (OX2R) | 173Yb | 1/100 |
| MHC class II (I-A/I-E) | Standard Biotools | 3174003B | Monoclone (M5/114.15.2) | 174Yb | 1/100 |
| CD18 | Biolegend | 101402 | Monoclone (M18/2) | 175Lu | 1/800 |
| CD73 | Biolegend | 127202 | Monoclone (TY/11.8) | 176Yb | 1/100 |

**Supplementary Table 2.** Antibodies used in imaging mass cytometry study.

| Antigen name | Source | Catalog number | Type | Metal tag | Dilution |
|---|---|---|---|---|---|
| CD45 | Standard Biotools | 3089005B | Monoclone (30-F11) | 89Y | 1/60 |
| CD13 | abcam | ab196576 | Monoclone (epr4059) | 115In | 1/170 |
| TNFa | Standard Biotools | 3141013B | Monoclone (MP6-XT22) | 141Pr | 1/90 |
| CD11c | Standard Biotools | 3142003B | Monoclone (N418) | 142Nd | 1/55 |
| GFAP | Standard Biotools | 3143030D | Monoclone (GA5) | 143Nd | 1/225 |
| CD68 | abcam | ab227458 | Monoclone (epr20545) | 144Nd | 1/65 |
| NeuN | abcam | ab209898 | Monoclone (epr12763) | 145Nd | 1/70 |
| Il10 | Biolegend | 505029 | Monoclone (JES5-16E3) | 146Nd | 1/60 |
| Vcam1 | ThermoFisher | PA5-47029 | Polyclone | 147Sm | 1/90 |
| Iba1 | abcam | ab221790 | Monoclone (epr16589) | 148Nd | 1/75 |
| Il12p40 | BD Biosciences | 554477 | Monoclone (C15.6) | 149Sm | 1/70 |
| Tau | CellSignaling | 29384SF | Monoclne (D1M9X) | 150Nd | 1/500 |
| gp91phox | BD Biosciences | 611415 | Monoclone (53/gp91[phox]) | 152Sm | 1/80 |
| CD274 | Standard Biotools | 3153016C | Monoclone (10F.9G2) | 153Eu | 1/60 |
| CD11b | Standard Biotools | 3154006B | Monoclone (M1/70) | 154Sm | 1/70 |
| CD40 | Biolegend | 102902 | Monoclone (HM40-3) | 155Gd | 1/100 |
| CD14 | Standard Biotools | 3156009B | Monoclone (Sa14-2) | 156Gd | 1/60 |
| TGFb | BD Biosciences | 555052 | Monoclone (A75-2) | 158Gd | 1/70 |
| F4/80 | Standard Biotools | 3159009B | Monoclone (BM8) | 159Tb | 1/65 |
| CD35 | BD Bioscences | 558768 | Monoclone (8C12) | 160Gd | 1/70 |
| pTau | ThermoFisher | MN1020 | Monoclone (AT8) | 161Dy | 1/60 |
| CD44 | Biolegend | 103002 | Monoclone (IM7) | 162Dy | 1/60 |
| CD172a | BD Biosciences | 552371 | Monoclone (P84) | 163Dy | 1/60 |
| CD164 (CX3CR1) | Standard Biotools | 3164023B | Monoclone (SA011F11) | 164Dy | 1/60 |
| MERTK | Biolegend | 151502 | Monoclone (2B10C42) | 165Ho | 1/60 |
| Arg1 | BD Biosciences | 610708 | Monoclone (Clone 19) | 166Er | 1/100 |
| GLUT1 | abcam | ab252403 | Monoclone (epr3915) | 167Er | 1/525 |
| CD206 | Biolegend | 141702 | Monoclone (C068C2) | 168Er | 1/60 |
| gingipain RgpB | University of Georgia, Athens, USA | Custom | Monoclone (25G3) | 169Tm | 1/70 |
| SiglecF | BD Bioscences | 552125 | Monoclone (E50-2440) | 170Er | 1/60 |
| Cox2 | CellSignaling | 73315SF | Monoclone (D5H5) | 171Yb | 1/125 |
| CD86 | Standard Biotools | 3172016B | Monoclone (GL1) | 172Yb | 1/50 |
| Amyloid | Biolegend | 800709 | Monoclone (4G8) | 173Yb | 1/80 |
| MHCII | Standard Biotools | 3174003B | Monoclone (M5/114.15.2) | 174Yb | 1/60 |
| CD18 | Biolegend | 101402 | Monoclone (M18/2) | 175 Lu | 1/60 |

| | | | | | |
|---|---|---|---|---|---|
| CCD73 | Biolegend | 127202 | Monoclone (TY/11.8) | 176Yb | 1/55 |
| Ly6G | Standard Biotools | 92J011195 | Monoclone (18A) | 195Pt | 1/55 |

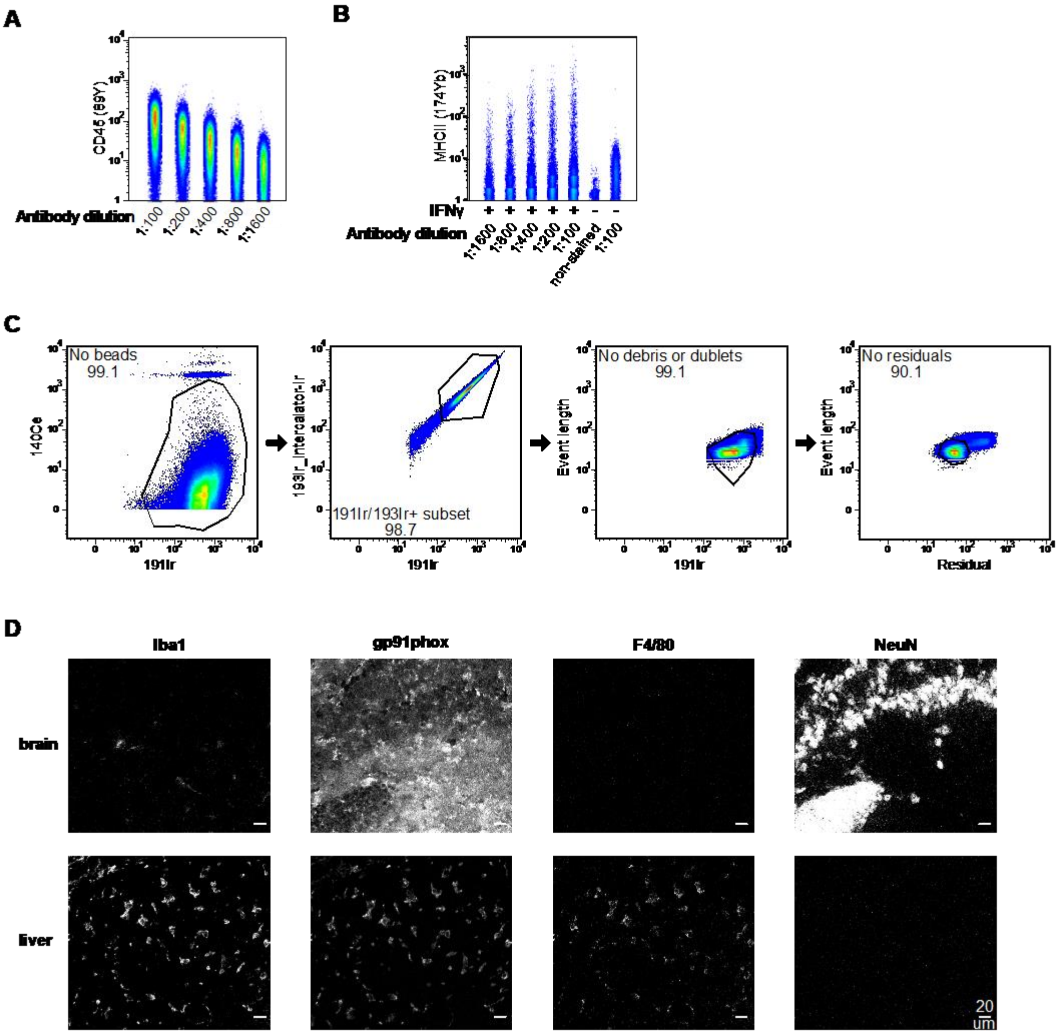


**Supplementary Figure 1. Antibody optimisation and quality-control workflow for CyTOF and imaging mass cytometry.**

**A.** Titration curve for anti-CD45 on fixed SIM-A9 cells. Serial dilutions (1:100–1:1,600) were stained for 30 min at 25 °C; the 1:400 dilution was selected for the main panel as it lies within the linear dynamic range while preserving signal-to-noise.

**B.** Titration of anti-MHC II on SIM-A9 cells pre-stimulated with 100 ng/ml IFN-γ for 16 h. The 1:100 dilution provided sufficient separation from the unstained control. **C.** Sequential gating strategy used for all CyTOF datasets: (1) removal of EQ bead events (140Ce); (2) selection of Intercalator-positive singlets (Ir191 vs Ir193, event length); (3) exclusion of debris and doublets; (4) residual events - non-cellular artifacts remaining after barcode deconvolution—were further excluded. See Methods and Extended Data Fig. 1 for subsequent downstream clustering. **D.** Antibody validation for imaging mass cytometry. Brain cryosections (top) and liver cryosections (bottom) were stained with anti-Iba1, gp91phox, F4/80 and NeuN (dilutions 1:50–1:200, 16 h at 4°C). Each antibody shows the expected regional distribution - microglia (Iba1, gp91phox, F4/80) and neuronal nuclei (NeuN) in hippocampus - and negligible NeuN signal in Kupffer cell–rich liver, confirming specificity. All images are single-channel, linearly scaled; scale bar, 20 µm. Detailed titration parameters and antibody lists are provided in Supplementary Tables 1 and 2.

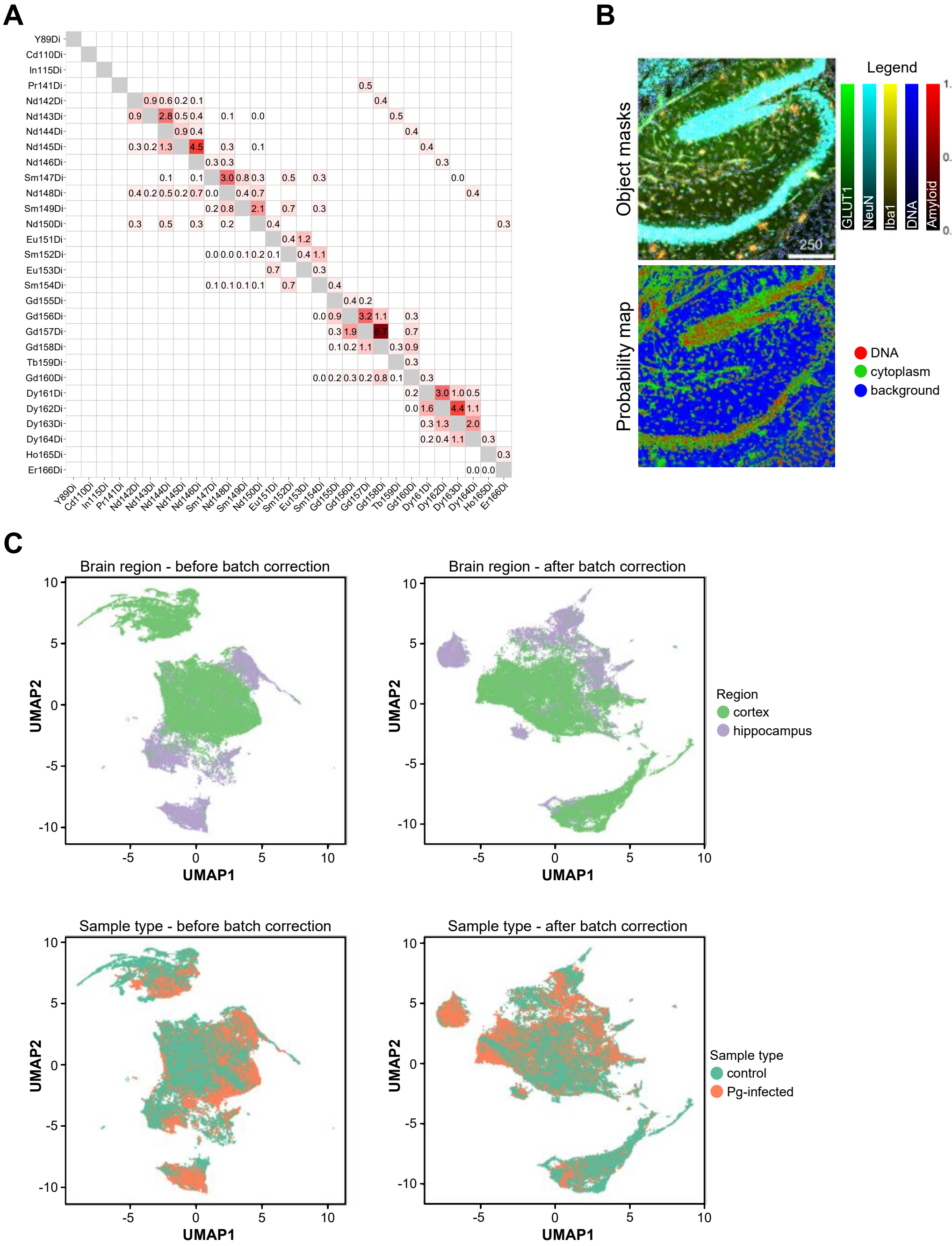


**Supplementary Figure 2. Quality control and data preprocessing of IMC dataset.**

**A.** Heat map of custom spillover compensation matrix generated from antibody microarray, showing signal intensities across channels used in the IMC panel. Each row represents a primary antibody–metal channel pair, and each column indicates measured signal in secondary channels. Spillover correction was applied to all channels except DNA intercalators (Ir191/193). **B.** Representative field of view from the hippocampus showing segmentation output. Top panel: object boundaries (white) overlaid on color-coded expression of GLUT1 (green), NeuN (cyan), Iba1 (yello), DNA (blue), and amyloid (red). Bottom panel: pixel classification probability maps from Ilastik, trained to identify nuclear (DNA), cytoplasmic, and background compartments. **C.** UMAP representations of segmented single-cell events before (left) and after (right) Seurat-based batch correction. Pre-integration plots are colored by brain region (top) and experimental group (bottom), highlighting slide-related variability. Post-correction UMAPs show improved integration across batches and preservation of biological separation by region or condition. Data reflect integrated hippocampal and cortical scans from PBS-treated control (n = 4) and P. gingivalis–infected (n = 3) hTau × 5×FAD mice. Methods for spillover compensation, pixel classification, and batch correction are detailed in the Methods section.

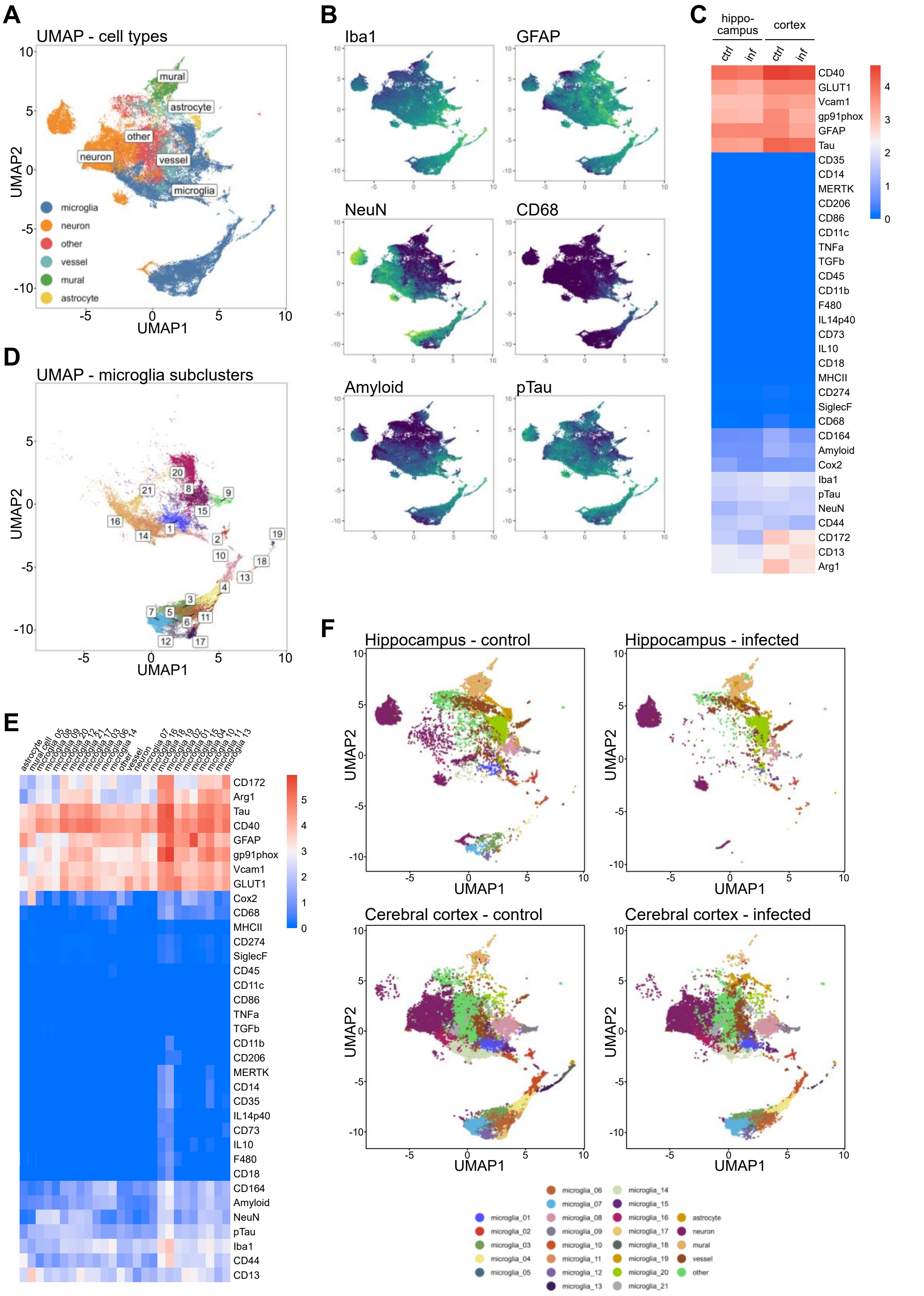


**Supplementary Figure 3. Analytical workflow and visualization of imaging mass cytometry dataset. A.** UMAP projection of all segmented cell-like objects (**n** = 78,622), colored by assigned major cell type based on marker expression (neurons, microglia, astrocytes, mural cells, blood vessels, and unclassified). **B.** UMAP overlays of normalized expression for representative antigens: Iba1 (microglia), GFAP (astrocytes), NeuN (neurons), CD68 (macrophages), amyloid, and phosphorylated Tau (pTau), illustrating regional and cell type–

specific distribution. **C.** Heat map of relative antigen expression across all samples, grouped by brain region (hippocampus or cortex) and experimental condition (control or infected). Only validated antibodies were included. **D.** UMAP of microglia-like objects (subset of a), clustered into 21 subpopulations using Rphenograph (k = 60) based on marker expression. **E.** Heat map of relative expression for all validated antigens across all idenfitied clusters with microglia subclustering. **F.** UMAPs of microglial subclusters separated by brain region and infection status: hippocampus–control, hippocampus–infected, cortex–control, and cortex–infected, showing differential cluster composition between experimental groups.

Dataset reflects cortical and hippocampal scans from hTau × 5×FAD mice exposed to *P. gingivalis* (infected, **n** = 3) or PBS (control, **n** = 4). Processing, segmentation, normalization, clustering, and batch correction were performed as described in the Methods.

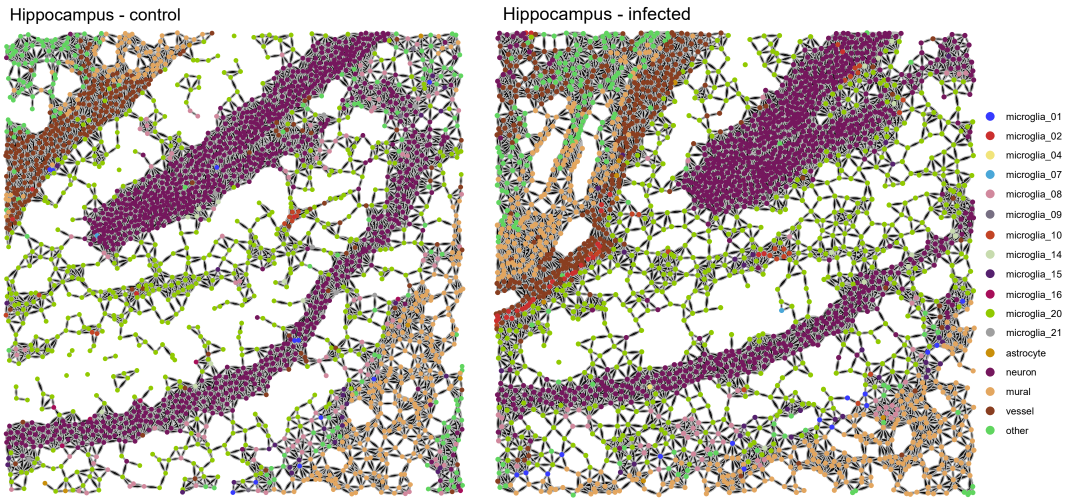


**Supplementary Figure 4. Cell-type–enriched region expansion maps for representative hippocampal ROIs.**

Representative regions from control and infected mice shown in Figure 4b are visualized here with cell-type–enriched expansions. Individual objects were assigned to one of six major cell types (neurons, microglia, astrocytes, endothelium, mural cells, or undefined) based on initial clustering. Each region was expanded radially to a 40-pixel radius to generate discrete local tissue domains. Colors represent dominant cell-type identity per region. This segmentation enabled downstream identification of spatial contexts (SCs) shown in the main figure.

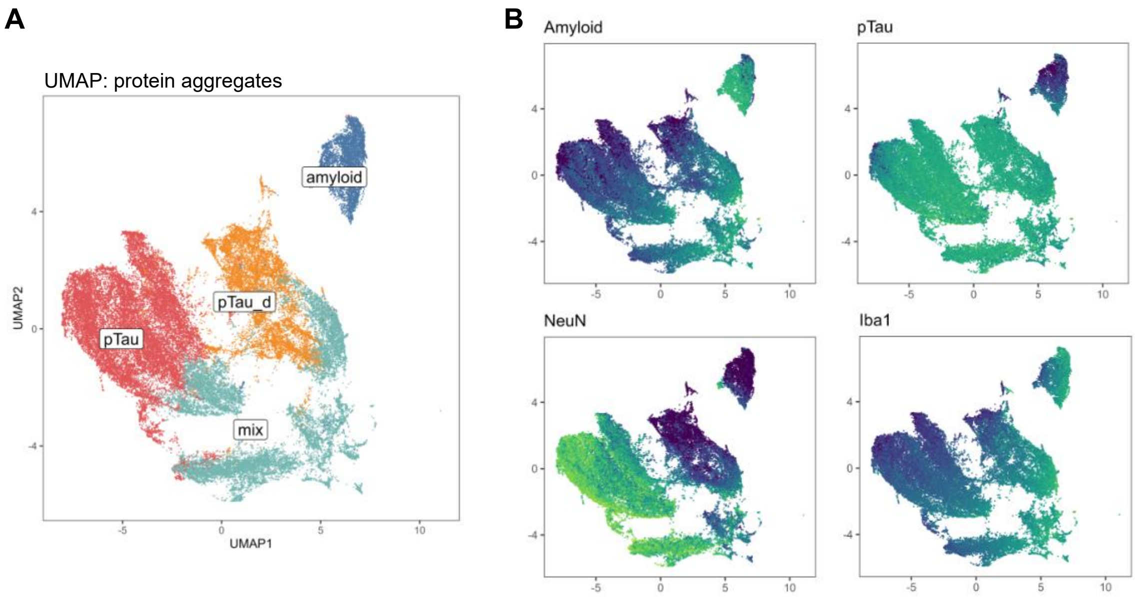


**Supplementary Figure 5. Protein aggregate characterization in hTau-knock-in x 5xFAD mice post-infection.**

**A.** Harmony-corrected UMAP visualization of protein aggregate objects segmented from imaging mass cytometry data, showing distinct clusters based on amyloid and pTau expression profiles. **B.** UMAPs colored by representative antigen expression levels for amyloid, pTau, NeuN, and Iba1, illustrating the spatial distribution and marker specificity within aggregate populations.
